# Statistical analysis of larval postspiracular filament length reveals continuous variation in Bolivian *Anopheles pseudopunctipennis* (Diptera: Culicidae)

**DOI:** 10.1101/2025.11.27.690953

**Authors:** Frédéric Lardeux, Deysi Vasquez, Rosenka Lardeux, Claudia Aliaga, Lineth Garcia, Libia Torrez

## Abstract

*Anopheles pseudopunctipennis* is a neotropical malaria vector widely distributed from northern Argentina and Chile to the southern United States. At the larval stage, it is characterized by posterior-lateral caudal filaments, which vary markedly in length within the same samples in Bolivia, with some individuals displaying unusually long filaments. The coexistence of individuals with relatively long and relatively short filaments raises the question of whether at least two distinct populations could be differentiated based on caudal filament length. This study examined filament-length distributions in two Bolivian dry-valley populations, El Chaco and Mataral, to determine whether variation reflects distinct subpopulations or continuous phenotypic variation within a single population. Distributions deviated from normality, exhibiting moderate skewness and tail heaviness, and the Generalized Error Distribution provided the best statistical fit. Examination of outliers and a targeted analysis of the distribution tail using multiple complementary methods showed that extreme values did not form a discrete secondary cluster but rather represented the upper continuum of the trait range. The two sites showed broadly comparable distributions, consistent with similar environmental conditions. These results emphasize the importance of using appropriate distributional models for continuous traits, highlight the occurrence of rare extreme phenotypes within otherwise homogeneous populations, and provide a baseline for future studies on the ecological and genetic determinants of caudal filament variation in *An. pseudopunctipennis*.

## Introduction

*Anopheles pseudopunctipennis* (Theobald, 1901) is a neotropical mosquito species with a broad distribution, ranging from northern Argentina and Chile to the southern United States, and including several Caribbean islands (Wilkerson et al., 2021). It is predominantly associated with valleys and foothills in mountainous landscapes, where it serves as a major vector of human malaria. At the larval stage, *An. pseudopunctipennis* is uniquely characterized by the presence of posterior-lateral “tails” (caudal filaments) on the spiracular lobe of the eighth abdominal segment. In Bolivia, substantial variation in tail length has been observed among sympatric populations, raising the possibility of distinct subpopulations (Lardeux et al., 2009). The coexistence of both short- and long-caudal filament individuals within the same geographic areas may indicate heterogeneous subpopulations in the dry valleys. Understanding whether distinct groups exist within sympatric populations of *An. pseudopunctipennis* in Bolivia is important, as such differentiation could be associated with behavioral or ecological traits that influence malaria transmission and responses to environmental factors.

Several approaches can be used to distinguish potential subpopulations within *Anopheles* species, including genetic analyses (e.g., mitochondrial and nuclear markers) (Estrada et al., 1993; Gómez et al., 2013), cytogenetic studies (Coetzee et al., 1999), geometric morphometrics (Dantur et al., 2011), and investigations of ecological or behavioral traits (Naranjo et al., 2016). While these methods provide powerful tools to reveal cryptic diversity, the present study uses traditional morphometric analyses based on simple linear measurements, which are also a cost-effective and accessible preliminary approach.

In the present study, two populations from Bolivia’s dry valleys, Mataral and El Chaco, were examined to evaluate whether caudal filament length could serve as a morphological marker of population differentiation, and whether the populations consist of distinct subgroups or are largely homogeneous, with only some extreme individuals, providing insight into the potential structuring of sympatric populations.

## Materials and methods

### Study area

The study was carried out in two localities of the dry valleys of the eastern Andes of Bolivia: Mataral (18°35ꞌ54.54” S, 65°9ꞌ6.84 W, Alt. 1512 m) and El Chaco (18°44ꞌ27.78” S, 65°8ꞌ54.60”, Alt. 1665m), located ≍37 km apart along the road connecting the two major cities of Cochabamba and Sucre (**Figure 1**). Each locality has about 1 000 inhabitants living in ≍100 houses near the Chico River (El Chaco) and Novillero River (Mataral). The main economic activity is agriculture. The valleys have a hot, dry steppe climate, with annual temperatures ranging from 7°C to 24°C and mean annual rainfall of 500–700 mm (Montes de Oca, 2005). Vegetation is predominantly deciduous dry forest, dominated by Anacardiaceae, Asteraceae, Cactaceae, Leguminosae, and Verbenaceae (Navarro & Maldonado 2002).

**Figure 1.**
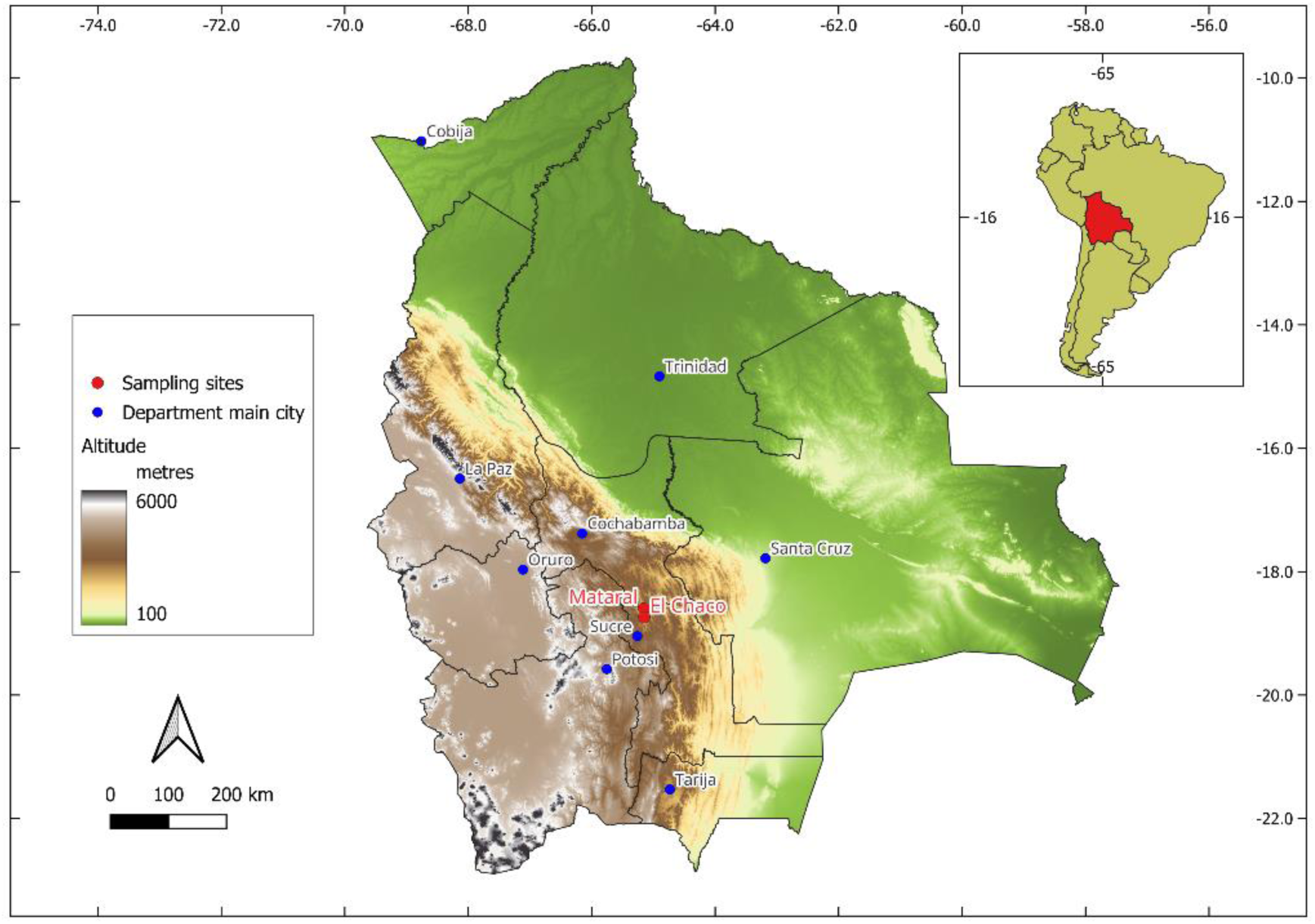
Geographic location of the two sampling sites, Mataral and El Chaco, in Bolivia

### Mosquito samples

*Anopheles pseudopunctipennis* larvae were collected in April 2019 from both sites using standard dipping techniques for mosquito larvae (O’Malley, 1995) in shallow pools along river margins containing clear, fresh water, exposed to sunlight, and with filamentous algae, which represent typical larval habitats for this species (Manguin et al., 1996). Collected fourth instar (L4) larvae were preserved in 70% ethanol for further analysis. At the study sites, only two *Anopheles* species co-occurred: *An. pseudopunctipennis* and *An. argyritarsis* (Lardeux et al., 2013). In the laboratory*, An. pseudopunctipennis* was distinguished from *An. argyritarsis* by the presence of the caudal filaments on the spiracular apparatus, which are absent in the latter and readily visible under a stereomicroscope. L4 larvae were clarified using 5% KOH, then rinsed in Marc-André solution, dehydrated through a graded ethanol series (70–100%), and finally mounted in Euparal (Karl Roth, Germany).

### Measurements of caudal filaments

Measurements were performed on digital images using *tpsUtil* and *tpsDig* softwares (Rohlf, 2015). Images were acquired with a Nikon D600 camera mounted on an Olympus CX-31 optical microscope. The lengths of the left and right caudal filaments were measured from the insertion point of seta XI of the spiracular apparatus to the distal end of the pigmented tail (**Figure 2**). All caudal filament and head images were calibrated with a 1000-*µ*m micrometer.

**Figure 2.**
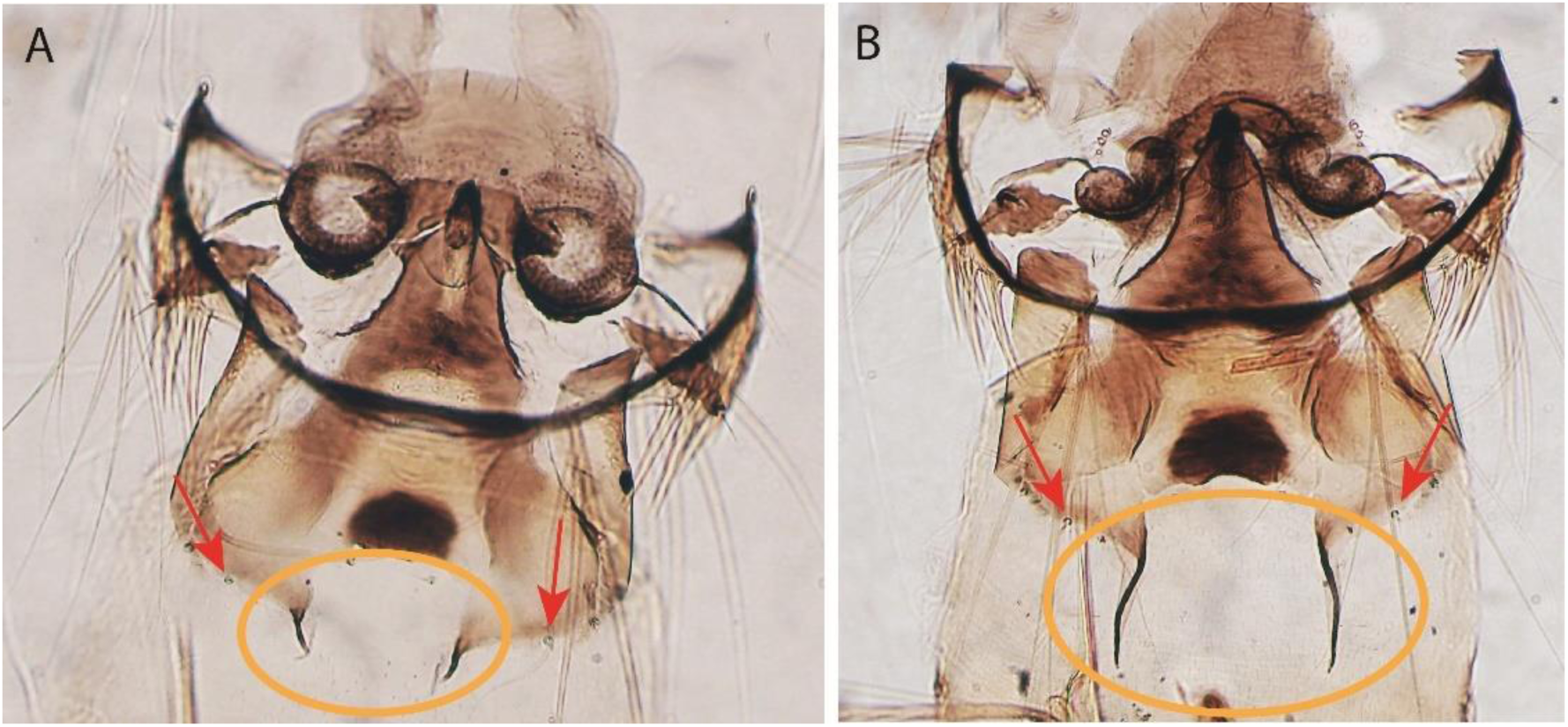
Spiracular apparatus of *An. pseudopunctipennis* showing the left and right filaments in orange ellipses. (A) Short filaments and (B) long filaments. Red arrows indicate the insertion point of setae XI, from which length measurements were taken down to the end of the black “tail”

### Statistical analysis

The analysis was conducted in the R statistical environment (version 4.3.0; R Core Team (2023)) using a comprehensive set of eight annotated R-scripts (*colas_1.R* to *colas_7.R*), available on Zenodo repository together with the datasets used in this study (Lardeux et al., 2025).

#### Data preparation and allometric adjustment

After verifying measurement accuracy, the mean caudal filament length (average to left and right) was calculated for each larva (R-script: *colas_1-mean.R* (Lardeux et al., 2025))

To account for potential allometric effects in caudal filament length measurements and for variation in overall larval size, the mean caudal filament length was standardized using the larval head-collar length, a morphological feature indicative of total larval size. Several transformations were applied to the data, including simple ratio (filament length / collar length), log-transformed ratio, residuals from ordinary least squares (OLS) linear regression (LM) and robust linear regression (RLM), residuals from log-transformed regressions, residuals from generalized additive models (GAM) (Lleonart et al., 2000), and values adjusted using standardized major axis (SMA) / major axis (MA) regression (Warton et al., 2006) (R-script: *colas_2_allometry.R* (Lardeux et al., 2025)). The effectiveness of each transformation was assessed using the absolute correlation between the transformed trait and head size, with lower absolute correlation indicating a better removal of allometric effects. The LM residuals effectively removed the allometric effect in both datasets (Mataral: R² = 7.42×10⁻¹⁶; El Chaco: R² = 6.24×10⁻¹⁶). Consequently, these allometric residuals (LM transformation) were used as size-corrected variables in subsequent analyses. Other transformations—ratio, log-ratio, SMA/MA-adjusted values, and residuals from GAM or RLM—showed higher correlations, meaning that the transformed variable still depended on head-collar size and was therefore less effective (**Table 1**). This approach ensures that subsequent analyses of morphometric variation are largely independent of overall body size, thereby isolating biologically meaningful variation in caudal filament length.

**Table 1.** Transformations tested for the removal of allometric effects. The metric corresponds to the Pearson correlation coefficient between the transformed variable and head collar size, except for the MA/SMA transformation, where it represents the coefficient of determination (R²) from the log–log regression. Rank for selection of the best transformation is based on the absolute value of the metric

| Transformation | Mataral |  | El Chaco |  |
| --- | --- | --- | --- | --- |
|  | Metric | Rank | Metric | Rank |
| Log(length) / Log(head) | -3.68E-01 | 8 | -3.29E-01 | 8 |
| MA / SMA | 3.23E-03 | 4 | 2.31E-02 | 6 |
| Length / Head | -3.68E-01 | 7 | -3.28E-01 | 7 |
| Residuals GAM | 1.81E-13 | 2 | -1.02E-15 | 2 |
| Residuals LM | -7.42E-16 | 1 | -1.91E-16 | 1 |
| Residuals Log(LM) | -9.27E-04 | 3 | 5.24E-03 | 3 |
| Residuals log(RLM) | 5.34E-03 | 5 | 1.19E-02 | 4 |
| Residuals RLM | 6.20E-03 | 6 | 1.64E-02 | 5 |

Abbreviations. MA / SMA: Major Axis / Standardized Major Axis regression adjustment; Residuals GAM: residuals from a Generalized Additive Model (non-linear adjustment); Residuals LM: residuals from linear regression (length of filament ~ head collar length); Residuals Log(LM): residuals from linear regression on log-transformed data. Residuals log(RLM): Residuals from robust linear regression on log-transformed data; residuals RLM: Residuals from robust linear regression (untransformed)

#### Inter-site comparison

Comparisons of sites were performed using the Kolmogorov–Smirnov (KS) test, which evaluates the maximum discrepancy between empirical cumulative distributions, and the Anderson–Darling test, which places additional weight on differences in the tails. Distributional shape was further assessed using bootstrap tests for skewness and kurtosis, where the test statistics correspond to the observed differences between sites and are evaluated via resampling (R-script: *colas_3_comparison_sites.R* (Lardeux et al., 2025)). These complementary approaches were used to determine whether data from El Chaco and Mataral could be legitimately combined or required separate analysis.

#### Descriptive statistics and distribution fitting

For each site (Mataral and El Chaco), the data set was summarized using standard descriptive statistics, including mean, median, standard deviation, variance, median absolute deviation (MAD), coefficient of variation, skewness, kurtosis, interquartile range (IQR), maximum and minimum values, and selected percentiles (5th, 25th, 75th, 95th) (R-script: *colas_4_stats.R* (Lardeux et al., 2025)). To assess the underlying distributional form, several distributions were fitted to each data set, including (1) univariate Generalized Error distribution (GED), Skew-*t,* Student’s *t*, Skew-normal, Normal, Log-Normal, Weibull, Gamma and Pareto, and (2) mixtures of Normal, skew-normal and Student’s *t* with two components to assess the presence of potential subpopulations (R-script: *colas_5_distrib_fit.R* (Lardeux et al., 2025)). The GED and the Student-*t* distribution extend the Gaussian family by accommodating excess kurtosis, thereby providing heavier tails and greater robustness to outliers. Skewed extensions (e.g., skew-normal, skew-*t*) add a shape parameter that accounts for distributional asymmetry, enabling flexible modeling of right- or left-skewed tail behavior. Model fit was assessed using Akaike Information criterion (AIC) and Bayesian Information Criterion (BIC), with model ranked accordingly.

As the hypothesis involved two distinct populations, the data were analyzed using three complementary approaches. First, a Bhattacharyya analysis (Bhattacharya, 1967) was performed to provide an initial assessment of potential subpopulation structure. These results were then used to initialize a two-component Gaussian Mixture Model (GMM) (hereafter referred to as GMM_seeded_), allowing the Expectation–Maximization (EM) algorithm to refine parameter estimates based on the Bhattacharyya-derived seeds. In parallel, a GMM with free parameters (hereafter referred to as GMM_free_) was fitted independently of any prior information to assess the robustness of the clustering. In both GMM approaches, multiple random starting points were used in the EM algorithm to evaluate the stability of parameter estimates. Component separation was quantified using Bhattacharyya distances (Bhattacharyya, 1946) and weighted and unweighted overlap coefficients (OVL) (Nowakowska et al., 2014), with larger distances and smaller OVL values indicating better discrimination between components. Posterior probabilities were computed for each observation to assign it to the most likely component, and observations assigned to the minor component were examined to determine whether they represented a rare tail of the distribution rather than a distinct subpopulation (R-script: *colas_6_GMM.R* (Lardeux et al., 2025).

#### Outlier detection methods

Outliers were identified using complementary classical and robust statistical approaches (R-script: *colas_7_outliers.R* (Lardeux et al., 2025)). Classical methods included the Interquartile Range (IQR) rule (Tukey, 1977), which flags values lying more than 1.5 × IQR beyond the first or third quartile, and the Z-score method (Barnett & Lewis, 1994), which identifies observations several standard deviations away from the mean under the assumption of approximate normality. Robust alternatives comprised the Median Absolute Deviation (MAD) criterion (Leys et al., 2013), flagging values exceeding typically three MADs from the median, and the Hampel identifier (Hampel, 1974), a moving-median variant of the MAD filter. The Grubbs test (Grubbs, 1950), applied iteratively, and the Generalized Extreme Studentized Deviate (ESD) test (Rosner, 1983), were used to detect one or several outliers under normality assumptions. To accommodate skewed distributions, the MedCouple-adjusted boxplot (Hubert & Vandervieren, 2008), was applied, which adapts boxplot fences according to sample skewness. Finally, model-based procedures included the Generalized Error Distribution (GED) approach (Nelson, 1991), identifying low-probability observations under a fitted GED model, and a two-component Gaussian Mixture Model (McLachlan & Peel, 2000), which classifies observations with low posterior probability of belonging to either main component as potential outliers. This combination of methods enhances both robustness and sensitivity across diverse data structures.

#### Tail Analysis of filament length distribution

To investigate the extreme values of larval filament lengths, a comprehensive tail analysis of the distribution was carried out, combining non-parametric, semi-parametric, and parametric approaches allowing robust characterization of both subpopulation structure and tail behavior (R-script: *colas_8_tail_analysis.R* (Lardeux et al., 2025)). Subpopulation structure and multimodality of the distribution was tested using the Dip test (Hartigan & Hartigan, 1985), kernel density estimation, the Silverman test (Silverman, 2018), the Excess Mass test (Müller & Sawitzki, 1991), Mode Tree analysis (Minnotte & Scott, 1993), Gaussian mixture modeling via BIC comparison (Fraley & Raftery, 2002), and the gap statistic (Tibshirani et al., 2001).

Threshold selection for peaks-over-threshold (POT) analysis was guided by mean residual life (mean-excess) plots (Embrechts et al., 1997) and adaptive quantile selection ensuring a sufficient number of exceedances (>20). Generalized Pareto distributions (GPD) were fitted to threshold exceedances using maximum likelihood estimation (Pickands, 1975; Coles, 2001), providing tail shape (*ξ*) and scale (*β*) parameters. Tail stability was assessed across multiple thresholds.

Complementary tail indices were estimated using the Hill estimator (Hill, 1975) and non-parametric estimators (Pickands, 1975; Dekkers et al., 1989) to validate tail heaviness. Block maxima were analyzed under the Generalized Extreme Value (GEV) distribution framework (Fisher & Tippett, 1928; Coles, 2001) with blocks of size chosen to balance sample resolution and reliability. Finally, tail conditional expectations (TCE) (Acerbi & Tasche, 2002) were computed for the 90th–99th percentiles.

## Results

### Inter-site comparison

The results of statistical tests comparing the distributions of the size-corrected variable between El Chaco and Mataral are presented in **Table 2** and **Figure 3**.

**Figure 3.**
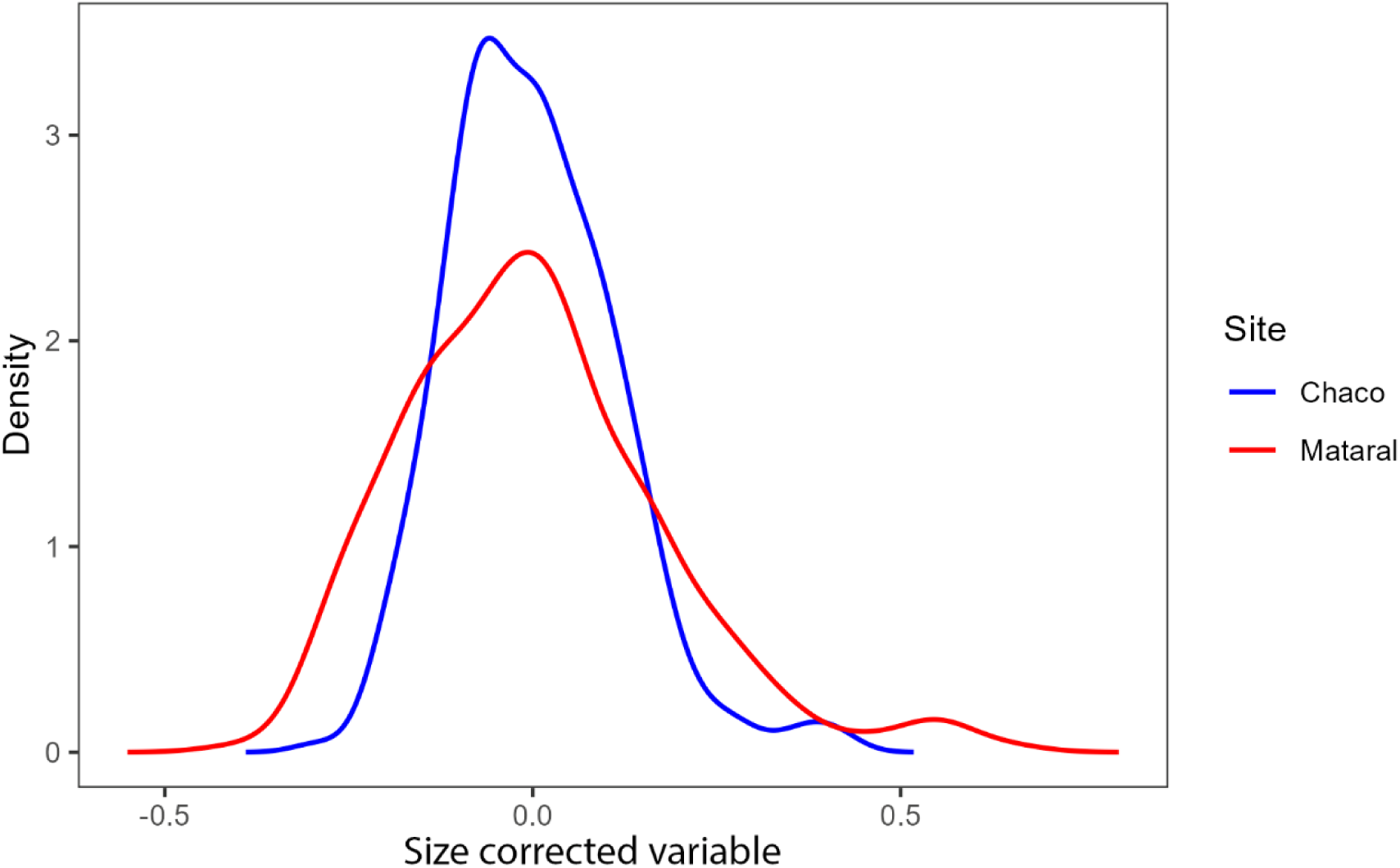
Kernel density estimates of the size corrected variable for El chaco (blue line) and Mataral (red line)

**Table 2.** Comparison of size-corrected variable (z-scored) between El Chaco and Mataral. The table shows the results of robust statistical tests: Kolmogorov–Smirnov for overall distribution, Mann–Whitney for median differences, and Levene test for homogeneity of variance. All tests indicate no significant differences between the two sites, suggesting comparable morphological patterns across locations

| Test | Statistics value | $p$ value | Significant | Interpretation |
| --- | --- | --- | --- | --- |
| Kolmogorov-Smirnov (raw data) | $D = 0.062$ | 0.001 | TRUE | Distributions different |
| Anderson–Darling | 6.93 | 0.0003 | TRUE | Distributions different |
| Skewness bootstrap | 0.134 | 0.6376 | FALSE | No significant difference |
| Kurtosis bootstrap | 0.0799 | 0.9096 | FALSE | No significant difference |
| KS (z-scores per site) | D = 0.0507 | 0.866 | FALSE | No significant difference |
| KS (global z-scores) | D = 0.164 | 0.001 | TRUE | Distributions different |

KS tests on raw and global z-scores detect overall distributional differences, including mean, variance, and shape. Anderson–Darling is sensitive to tail differences. Skewness and kurtosis bootstrap tests assess differences in asymmetry and tail heaviness. KS on site-standardized z-scores isolates shape differences after normalization.

The two-sample Kolmogorov–Smirnov (KS) test on raw data indicated a modest but significant difference (D = 0.164, *p* = 0.0011), suggesting that the empirical cumulative distributions are not identical. Similarly, the Anderson–Darling (AD) k-sample test, which is particularly sensitive to differences in the tails, was highly significant (AD ≈ 6.93, p ≈ 0.00032). In contrast, bootstrap tests for differences in skewness and kurtosis were non-significant (*p_skew* = 0.638; *p_kurt* = 0.910), indicating broadly comparable asymmetry and tail characteristics.

Kolmogorov–Smirnov tests on site-standardized z-scores (KS = 0.051, *p* > 0.05) suggest that the general distributional shapes are similar once differences in mean and variance are removed. However, the global z-score KS test remained significant (D = 0.164, *p* < 0.01), reflecting small but detectable overall differences in distributions when the data are combined. Descriptive statistics confirm broadly comparable central tendencies (means ≈ 0) and dispersions (SD = 0.114 and 0.177), with slightly right-skewed distributions and moderate kurtosis.

Taken together, these results indicate a common morphological pattern consistent with a single underlying population structure characterized by continuous variation and occasional large individuals. The subtle tail differences justify analyzing Chaco and Mataral separately, preserving the ability to detect site-specific deviations while acknowledging broadly homogeneous developmental and environmental conditions.

### The Mataral site

#### Basic characteristics of the distribution

The analysis was based on 263 individual measurements of larval filament length. The raw variable exhibited an average length of 1.096 mm (SD = 0.178), ranging from 0.685 mm to 1.749 mm (**Table 3**). After allometric correction, the size-corrected variable had a mean value close to zero (−2.9 × 10⁻¹⁷) and a variance of 0.031, consistent with successful normalization while preserving the dispersion of values (**Table 3**). The standard deviation of the transformed variable (0.177) and its interquartile range (0.228) indicated moderate dispersion around the mean.

**Table 3.** Basic characteristics of the distributions of datasets for Mataral and El Chaco sites. Datasets comprised of raw data (mean of left and right filament length) and transformed mean for allometric adjustment

| Variable | Mataral |  | El Chaco |  |
| --- | --- | --- | --- | --- |
|  | Raw data | Allometric | Raw data | Allometric |
| n | 263 | 263 | 297 | 297 |
| Mean | 1.096 | -2.88E-17 | 1.034 | -5.46E-18 |
| Variance | 0.032 | 0.031 | 0.013 | 0.013 |
| SD | 0.178 | 0.177 | 0.116 | 0.114 |
| IC95-mean | 0.021 | 0.021 | 0.013 | 0.013 |
| CV* | 0.16 | -6.16E+15 | 0.11 | -2.09E+16 |
| Median | 1.080 | -0.018 | 1.026 | -0.010 |
| Min | 0.685 | -0.400 | 0.738 | -0.293 |
| Max | 1.749 | 0.646 | 1.444 | 0.420 |
| Range | 1.064 | 1.046 | 0.706 | 0.713 |
| Max / Min | 2.554 | -1.614 | 1.957 | -1.436 |
| MAD | 0.115 | 0.114 | 0.079 | 0.078 |
| IQR | 0.222 | 0.228 | 0.158 | 0.151 |
| Q5 | 0.840 | -0.259 | 0.860 | -0.169 |
| Q25 | 0.964 | -0.129 | 0.948 | -0.077 |
| Q75 | 1.187 | 0.099 | 1.106 | 0.074 |
| Q95 | 1.400 | 0.307 | 1.229 | 0.182 |
| Q99 | 1.654 | 0.554 | 1.424 | 0.373 |
| Skewness | 0.799 | 0.800 | 0.694 | 0.665 |
| Kurtosis | 1.107 | 1.084 | 1.025 | 0.998 |

Abbreviations. MAD: Mean Absolute Deviation; IQR: interquartile range(*i.e.*, Q75 – Q25); $Q_x$ : $x^{\text{th}}$ percentile; IC95\_mean: 95% confidence interval half-width for the mean (*i.e.* the 95% confidence interval is [mean - IC95\_mean, mean + IC95\_mean]); CV: coefficient of variation

The range (−0.400 to 0.646) was asymmetrical, with the upper tail (0.646) extending farther from the median (−0.018) than the lower bound, suggesting the presence of a few individuals with unusually long filaments. The distribution was slightly right-skewed (skewness = 0.80) and moderately leptokurtic (kurtosis = 1.08), consistent with a near-normal shape but with heavier tails (**Figure 4**).

**Figure 4.**
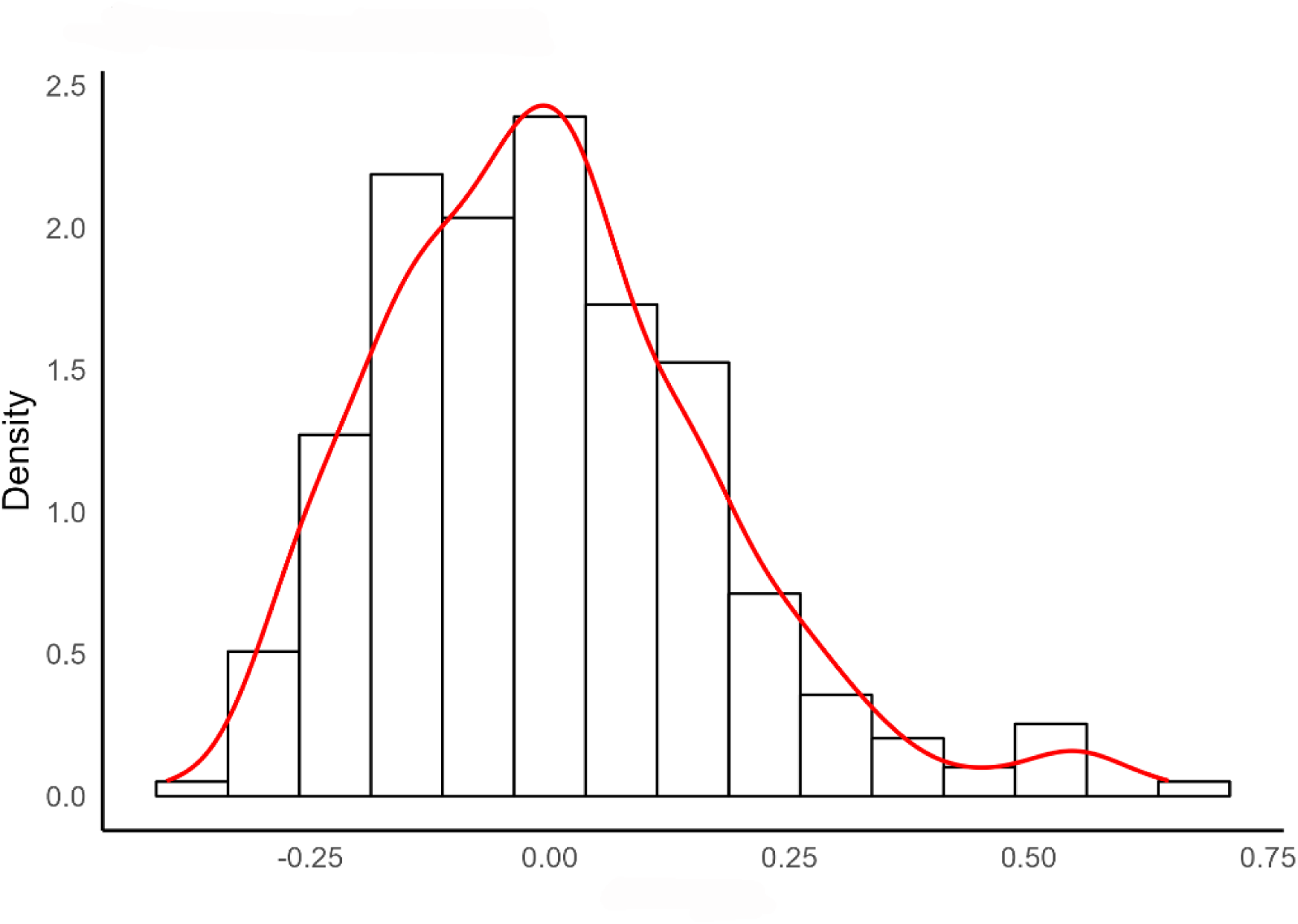
Histogram and density function of the Mataral dataset. The variable is the size-corrected variable and represents the allometric residuals (LM-transformed mean of right and left filament lengths relative to head-collar length)

The maximum value exceeded the absolute minimum by a factor of 1.61. The interquartile range (0.228) and median absolute deviation (0.114) suggested reasonable central clustering, although the gap between the 95th percentile (0.307) and the maximum (0.646) represented 32% of the total range, highlighting the extended right tail. Because the variable was centered around zero, the coefficient of variation (CV ≈ −6 × 10¹⁵) has no meaningful interpretation and was therefore not used as an indicator of relative dispersion.

#### Outlier detection

As shown in **Table 4**, the number of outliers detected in Mataral varied with the method applied but remained consistently low overall. Conservative approaches yielded few outliers: Grubbs’ iterative test detected only one observation (0.38%), while the ESD method identified none. Moderate methods, including Z-score, MedCouple, Hampel, and GED, consistently flagged 5–6 observations (1.9–2.3%). More liberal approaches identified more cases: the IQR method detected 7 outliers (2.7%), MAD identified 13 (4.9%), and the mixture-based approach flagged 14 observations (5.3%). This variation reflected differing sensitivity to tail behavior rather than algorithmic inconsistency.

**Table 4.** Detection of statistical outliers among larval postspiracular filament measurements of *An. pseudopunctipennis* from Mataral and El Chaco. Methods used, number of outliers detected and proportion of outliers in the whole sample

| Method | Mataral |  | El Chaco |  |
| --- | --- | --- | --- | --- |
|  | Number of outliers | Proportion of sample (%) | Number of outliers | Proportion of sample (%) |
| IQR | 7 | 2.7 | 5 | 1.7 |
| MAD | 13 | 4.9 | 10 | 3.4 |
| Z-score | 5 | 1.9 | 4 | 1.4 |
| Grubbs iterative | 1 | 0.4 | 2 | 0.7 |
| ESD | 0 | 0.0 | 0 | 0.0 |
| MedCouple | 5 | 1.9 | 5 | 1.7 |
| Hampel | 6 | 2.3 | 4 | 1.4 |
| GED | 6 | 2.3 | 5 | 1.7 |
| Mixture2G | 14 | 5.3 | 15 | 5.1 |

Abbreviations. MAD: Median Absolute Deviate; ESD: Generalized Extreme Studentized Deviate test; GED: Generalized Error Distribution-based detection method; Mixture 2G: the two-component Gaussian mixture model

The convergence of most methods around 5–6 outliers (approximately 2% of *n* = 263) suggested a modest but detectable subset of individuals with unusually long filaments. However, even under the most liberal criterion, only 14 observations were flagged, indicating a small potential subpopulation. No extremely short filaments were detected. These results supported the visual impression of a slightly extended right tail rather than a clearly distinct secondary group. This pattern is illustrated in **Figure 5**, which displays the individual filament lengths with IQR-defined outliers and the 95% and 99% GED-based thresholds. The plot confirms that only a few observations lie slightly beyond the upper limits, consistent with a light right-tail extension rather than a distinct subgroup.

**Figure 5.**
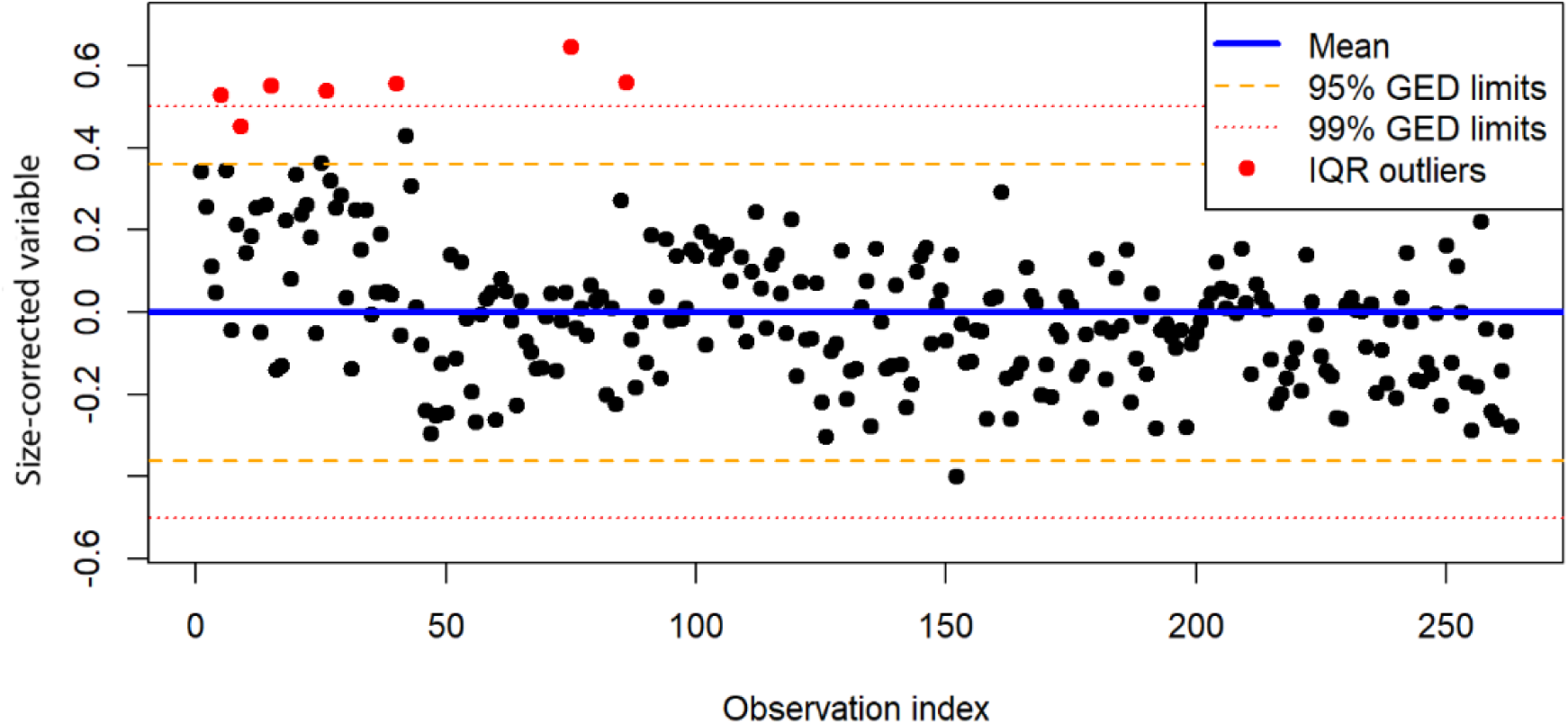
Distribution of individual data points from the Mataral dataset plotted by index (X-axis) and size-corrected variable (Y-axis). The solid horizontal line indicates the overall mean, while dashed lines represent the 95% and 99% GED (Generalized Error Distribution) limits. Red points denote outliers identified using the interquartile range (IQR) method

#### Distribution model fitting

**Table 5** summarizes the probability distributions tested for fitting the Mataral dataset, including both simple and two-component mixture models. Among the 14 tested models, the Generalized Error Distribution (GED) provided the best fit, with the lowest AIC (−327.0) and BIC (−316.3), followed by the skew-normal distribution (ΔAIC = 145.3) and the two-component Gaussian mixture (ΔAIC = 148.5).

**Table 5.** Comparison of statistical models fitted to the filament length residuals in *An. pseudopunctipennis* larvae from Mataral. Log-likelihood, Akaike Information Criterion (AIC), Bayesian Information Criterion (BIC), the differences relative to the best AIC (ΔAIC) and BIC (ΔBIC), and the ranking of each model based on ΔAIC and ΔBIC

| Model | Log Likelihood | AIC | BIC | ΔAIC | ΔBIC | Rank by ΔAIC | Rank by ΔBIC |
| --- | --- | --- | --- | --- | --- | --- | --- |
| GED | 166.51 | -327.01 | -316.29 | 0.00 | 0.00 | 1 | 1 |
| Skew Normal | 93.85 | -181.70 | -170.99 | 145.31 | 145.31 | 2 | 3 |
| Battacharya | 95.30 | -180.61 | -162.75 | 146.40 | 135.68 | 3 | 2 |
| GMMseeded ($k=2$) | 95.30 | -180.61 | -162.75 | 146.40 | 135.68 | 3 | 2 |
| GMMfree ($k=2$) | 94.27 | -178.53 | -160.67 | 148.48 | 155.62 | 5 | 5 |
| Mixture Skew-Normal ($k=2$) | 93.59 | -177.17 | -159.31 | 149.84 | 156.98 | 6 | 6 |
| Skew-$t$ | 91.14 | -174.27 | -159.98 | 152.74 | 156.31 | 7 | 7 |
| $t$-Student | 85.99 | -165.97 | -155.26 | 161.04 | 161.04 | 8 | 8 |
| Normal | 82.22 | -160.43 | -153.29 | 166.58 | 163.01 | 9 | 9 |
| Weibull | 78.48 | -152.96 | -145.81 | 174.05 | 170.48 | 10 | 10 |
| Mixture $t$-Student ($k=2$) | 78.18 | -142.36 | -117.36 | 184.65 | 198.94 | 11 | 11 |
| Gamma | 62.39 | -120.77 | -113.63 | 206.24 | 202.66 | 12 | 12 |
| Log-Normal | -70.52 | 145.05 | 152.19 | 472.06 | 468.49 | 13 | 13 |
| Pareto | -3318.98 | 6641.96 | 6649.10 | 6968.97 | 6965.40 | 14 | 14 |

GED: Generalized Error Distribution. GMM: Gaussian Mixture Models. All mixture models were fitted as 2 components mixtures ( $k=2$ ). The Battacharya model found 2 components

The fitted GED parameters (*μ* = 0.074, *σ* = 0.423, *ν* = 5.496) indicated a distribution slightly shifted toward positive values, with moderate spread and a shape parameter substantially exceeding two, suggesting lighter tails than Gaussian—consistent with the mild skewness and kurtosis previously observed. Classical normal and Student’s *t* models performed less well (ΔAIC > 160), indicating that deviations from Gaussianity were small but statistically detectable. Two-component mixture models provided only marginal improvement over single-component models. The Gaussian two-component analysis (GMM_free_, GMM_seeded_ and Battacharrya analysis) yielded slight increases in log-likelihood, whereas other mixture models (skew-Normal, Student-*t*, skew-*t*) performed worse (ΔAIC > 149 compared with the GED) providing no evidence for discrete subpopulations.

#### Two-component GMM specific analysis

##### Model fitting

The two seeding-based models (Bhattacharya and GMM-seeded) converged toward highly similar parameter estimates. However, the Bhattacharya method struggled to clearly identify the second component, which was very small and only marginally distinguishable in the data (**Figure 6**).

**Figure 6.**
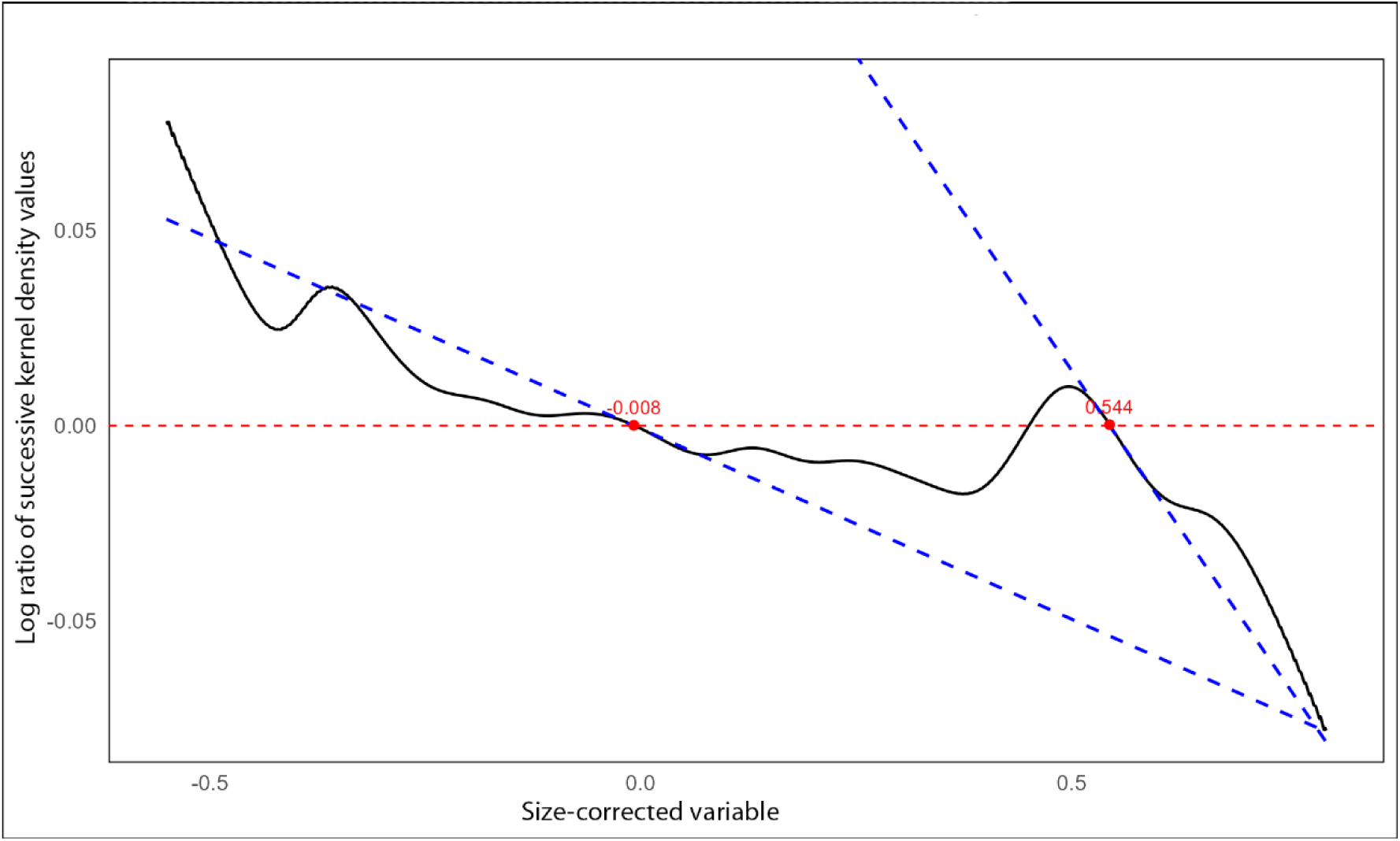
Bhattacharya plot of the Mataral dataset. The x-axis represents the size-corrected variable for filament lengths, and the y-axis shows the log ratio of successive kernel density values (Bhattacharya transformation). Approximate linear segments reveal the presence of two Gaussian components, from which means, variances, and proportions were estimated

The major component (component 1) had a mean around −0.015, variance ≈ 0.024, and accounted for ∼97 % of the data, whereas the minor component (component 2) had a mean ≈ 0.55, variance ≈ 0.003, and represented only ∼2.6 % of observations. The visual inspection of the reassigned data points and component histograms (**Figure 7**, **Figure 8**) further illustrates this difficulty in resolving the weak secondary mode.

**Figure 7.**
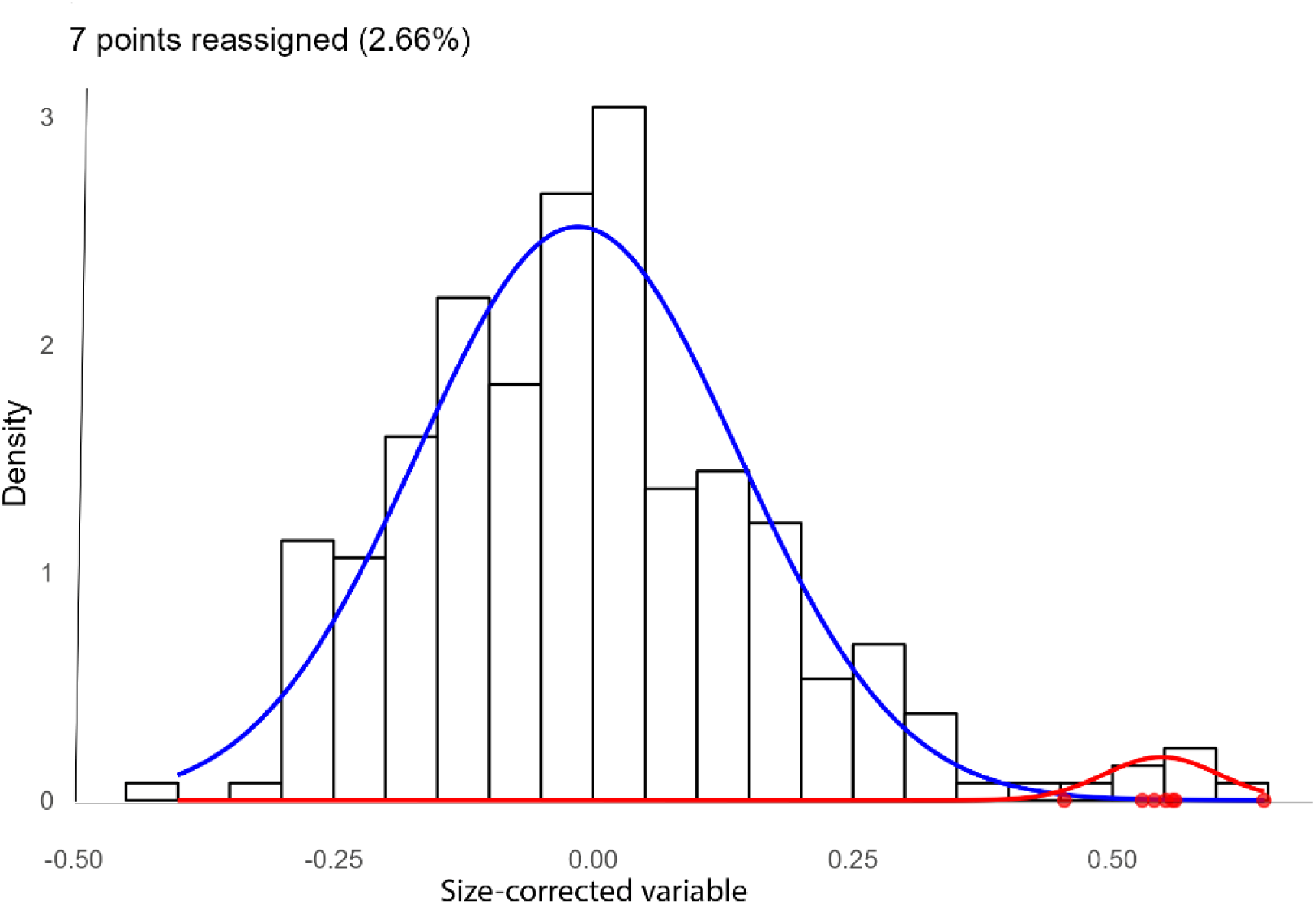
Histogram of the Mataral dataset showing fitted GMM_seeded_ components and reassigned data points by posterior probability

**Figure 8.**
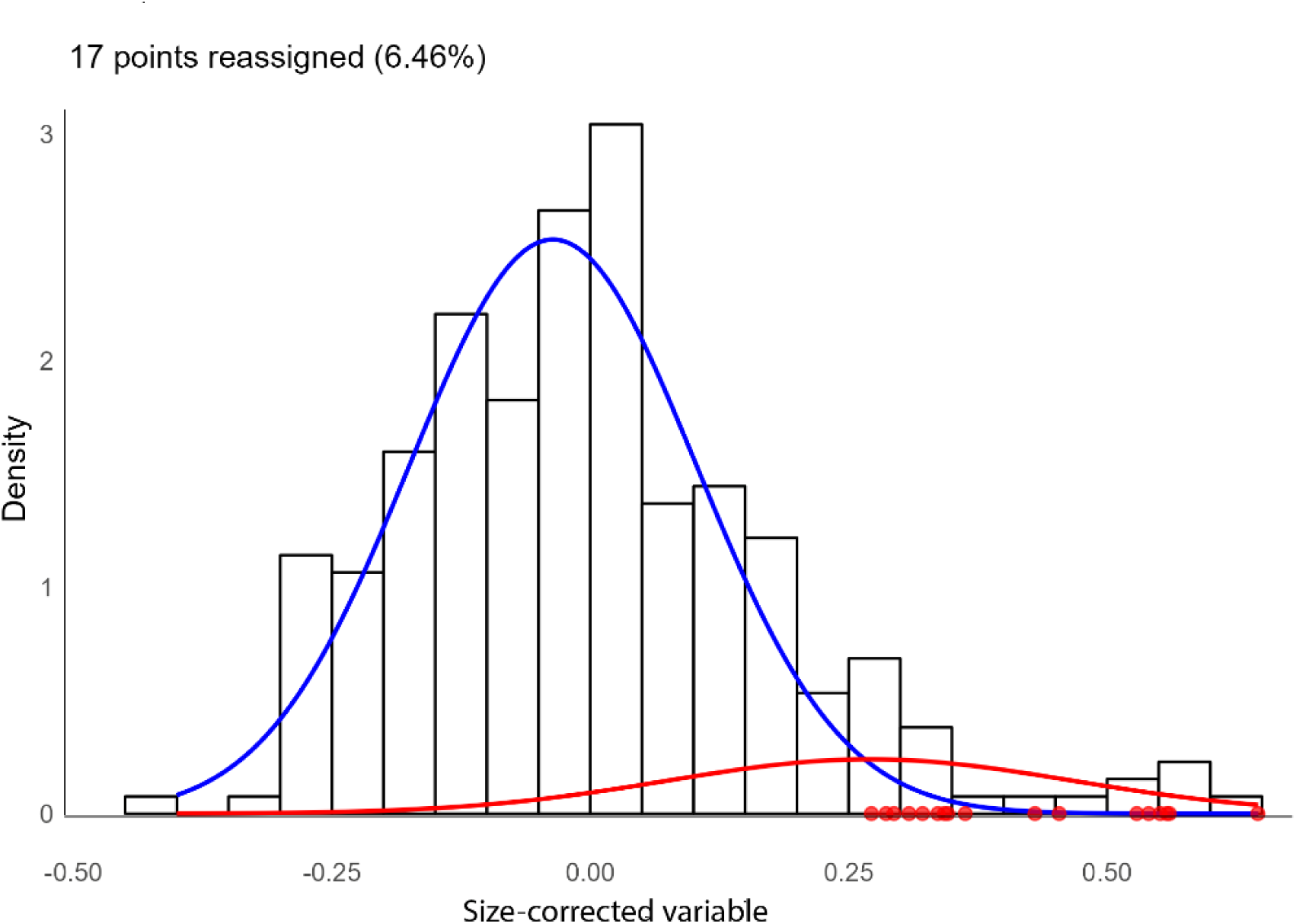
Histogram of the Mataral dataset showing fitted GMM_free_ components and reassigned data points by posterior probability

By contrast, the GMM_free_ estimated a larger minor component (component 2: mean = 0.271, variance = 0.038, proportion = 12%), while the major component (component 1: mean = −0.036, variance = 0.019, proportion = 88%) shifted slightly relative to the seeded models. The log-likelihood, AIC, and BIC values for GMM_free_ were slightly worse than those of the seeded models, reflecting a trade-off between model fit and stability (**Table 6).**

**Table 6.**
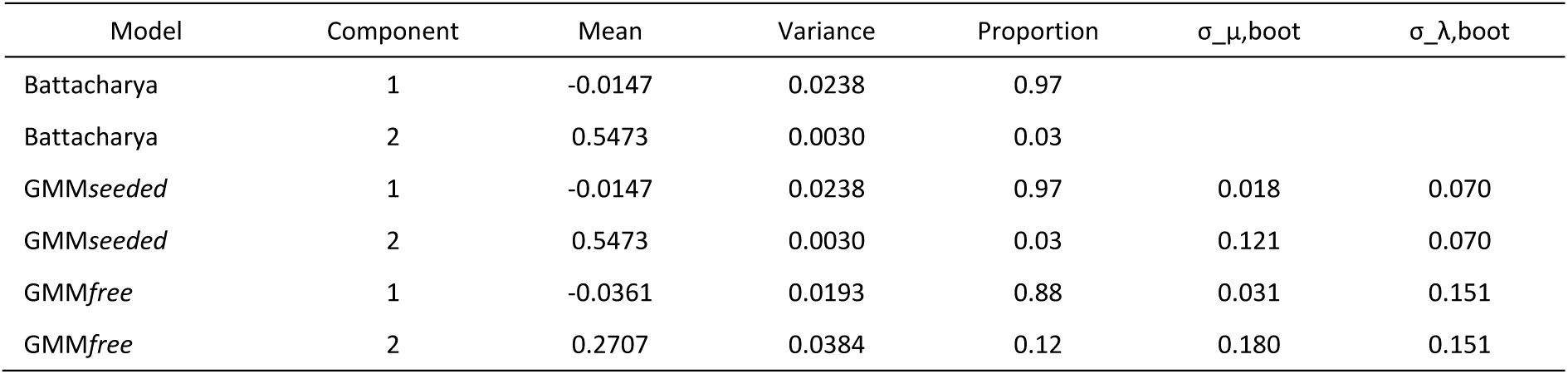
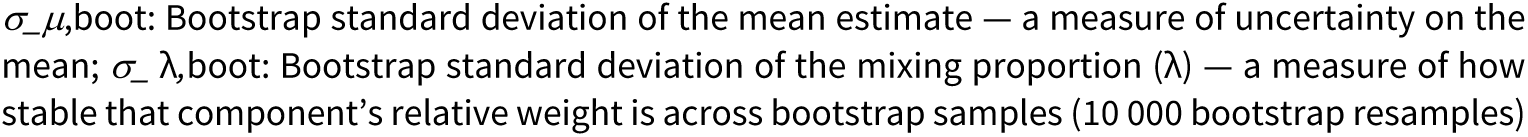
Summary of parameter estimates for the Mataral dataset obtained using three estimation approaches: Bhattacharya method, Gaussian mixture model (GMM) initialized with Bhattacharya estimates (“GMM-seeded”), and freely initialized GMM (“GMM-free”). For each component, mean, variance, and mixing proportion (λ) are reported, along with bootstrap standard deviations of the mean (σ₍μ,boot₎) and mixing proportion (σ₍λ,boot₎)

| Model | Component | Mean | Variance | Proportion | $\sigma_{\mu, \text{boot}}$ | $\sigma_{\lambda, \text{boot}}$ |
| --- | --- | --- | --- | --- | --- | --- |
| Battacharya | 1 | -0.0147 | 0.0238 | 0.97 |  |  |
| Battacharya | 2 | 0.5473 | 0.0030 | 0.03 |  |  |
| *GMMseeded* | 1 | -0.0147 | 0.0238 | 0.97 | 0.018 | 0.070 |
| *GMMseeded* | 2 | 0.5473 | 0.0030 | 0.03 | 0.121 | 0.070 |
| *GMMfree* | 1 | -0.0361 | 0.0193 | 0.88 | 0.031 | 0.151 |
| *GMMfree* | 2 | 0.2707 | 0.0384 | 0.12 | 0.180 | 0.151 |

$\sigma_{\mu,boot}$ : Bootstrap standard deviation of the mean estimate — a measure of uncertainty on the mean; $\sigma_{\lambda,boot}$ : Bootstrap standard deviation of the mixing proportion ( $\lambda$ ) — a measure of how stable that component's relative weight is across bootstrap samples (10 000 bootstrap resamples)

#### Component stability and separation

Component stability was assessed via bootstrapping (**Table 6**). In the GMM_seeded_ model, component 1 was highly stable (mean bootstrap SD = 0.018, *λ* bootstrap SD = 0.070), whereas component 2 was unstable, reflecting its very low proportion (2.6%) and limited data support. Similarly, in the GMM_free_ model, component 1 was stable (mean bootstrap SD = 0.031, *λ* bootstrap SD SD = 0.151), but component 2 was unstable (mean bootstrap SD = 0.180, *λ* bootstrap SD = 0.151), consistent with its minor representation (11.8%) and high bootstrap variability.

Distance metrics between components further highlighted differences among model (**Table 7**). Bhattacharyya and Mahalanobis distances were substantially larger in GMM_seeded_ model (3.17 and 4.85, respectively) than in the GMM_free_ model (0.44 and 1.81 respectively), suggesting that the free model components were closer in distributional space and that the minor component might be partially overlapping with the tail of the major component. Overlap analysis confirmed this: the minor component in the seeded models exhibited very low overlap with the major component (OVL_unweighted_ = 0.015, OVL_weighted_ = 0.002), whereas GMM_free_ showed substantial overlap (OVL_unweighted_ = 0.367, OVL_weighted_ = 0.071) (**Table 7**).

**Table 7.** Component separation and overlap for the Mataral dataset, with distances (Bhattacharyya, Mahalanobis) and OVL values indicating the distinctness of component distributions.

| Model | Bhattacharyya distance | Mahalanobis distance | OVL<sub>unweighted</sub> | OVL<sub>weighted</sub> |
| --- | --- | --- | --- | --- |
| Batttacharya | 3.17 | 4.85 | 0.015 | 0.002 |
| GMM<sub>free</sub> | 0.44 | 1.81 | 0.015 | 0.002 |
| GMM<sub>seeded</sub> | 3.17 | 4.85 | 0.367 | 0.071 |

#### Cross-model concordance

Cross-model stability analysis **(Table 8**) indicated high concordance between models (96–100%), although correlations in component assignment probabilities were higher between the seeded models than between the seeded and GMM_free_.

**Table 8.** Concordance of component assignments between models for the Mataral dataset, with percentage agreement and correlations of membership probabilities indicating similarity of clustering.

| Model 1 | Model 2 | Concordance | Corr-p1 | Corr-p2 |
| --- | --- | --- | --- | --- |
| Batttacharya | GMM<sub>free</sub> | 96.2 | 0.696 | 0.696 |
| Batttacharya | GMM<sub>seeded</sub> | 100 | 1 | 1 |
| GMM<sub>free</sub> | GMM<sub>seeded</sub> | 96.2 | 0.696 | 0.696 |

Concordance: percentage of individuals assigned to the same component in both models; Corr-p1 and Corr-p2: correlation of component membership probabilities for Model-1 and Model-2, respectively.

#### Posterior classification

Examination of reassigned points (Supplementary Table S1) showed that points assigned to the minor component in the seeded models were generally concentrated in the extreme right tail of the distribution. In GMM_free_, the minor component encompassed a broader range of values, including points closer to the main distribution. The GMM_seeded_ assigned seven observations to the minor component, with posterior probabilities of assignment to the minor component *P*(comp2) = 0.64–0.99), clustered around 0.53–0.65 near the second Gaussian mean (0.55 ± 0.06). One intermediate case (value = 0.454; *p* = 0.64) indicates a gradual transition. Given its low weight (2.6 %) and instability (**Table 6**), this component likely represents a small right-hand tail rather than a distinct mode (**Figure 7**).

In contrast, the GMM_free_ classified 16 points to the minor component (Table S1; weight = 11.8 %), spanning 0.27–0.65 with *P*(comp2) = 0.52–0.99. High-probability assignments (>0.95) occurred above 0.43, while lower values (0.27–0.36) overlapped with the main component (**Figure 8**). This reflects tail asymmetry rather than a separate mode, consistent with lower Bhattacharyya and Mahalanobis distances and higher overlap (**Table 8**).

Overall, the analysis of Gaussian mixture components indicates that the data are predominantly unimodal with a dominant Gaussian component and a small, unstable secondary component (<10 % of the data) likely representing a tail of the main distribution rather than a distinct subpopulation.

#### Tail-specific analysis

##### Multimodality and subpopulation tests

All three formal tests of multimodality—the Dip test (*p* = 0.48), Silverman’s test (*p* = 0.31), and the Excess Mass test (*p* = 0.09)—indicated no significant deviation from unimodality. Similarly, the mode tree analysis detected a single dominant mode. Model-based clustering yielded a minimal improvement in BIC (ΔBIC = 2), favoring a single-component Gaussian model. While the Gap statistic suggested *K*=2 as the optimal number of clusters (Gap = 0.196 vs 0.024 for K = 1; SE ≈ 0.107) (**Table 9**), the increase was moderate, indicating that the population remains largely homogeneous and the second cluster likely reflects subtle variation rather than a biologically distinct group (Tibshirani et al., 2001).

**Table 9.** Gap statistic and standard error (SE) for different numbers of clusters in the Mataral dataset.

| K<br>(number of<br>clusters) | Gap value | Standard error<br>(SE) |
| --- | --- | --- |
| 1 | 0.024 | 0.101 |
| 2 | 0.196 | 0.107 |
| 3 | 0.149 | 0.095 |

An examination of the sorted data revealed 14 unusually large gaps between consecutive values, indicating mild irregularities in the upper tail of the distribution; however, these gaps did not correspond to distinct clusters or separate subpopulations. Together, these results suggested that the dataset was globally unimodal but exhibited a few statistically distinct upper-tail observations rather than a clear secondary population.

#### Characterization of the right-hand tail

To quantify the behavior of the extreme upper tail, Generalized Pareto Distributions (GPD) were fitted above the 0.90 and 0.95 quantiles (thresholds 0.226 and 0.307) (**Table 10**). Both fits yielded negative shape parameters (ξ = −0.167 to −0.815), indicating a short or bounded tail rather than a heavy-tailed process. The scale parameter increased from 0.154 to 0.282 as the threshold rose, consistent with greater variability among the most extreme values. Standard errors were relatively large, especially for ξ (0.333 and 0.549), reflecting the limited number of exceedances (27 and 14 observations, respectively).

**Table 10.** Generalized Pareto Distribution (GPD) fits to the upper tail of the Mataral dataset.

| Quantile | Threshold | Exceedances | Scale | Shape | SE Scale | SE Shape |
| --- | --- | --- | --- | --- | --- | --- |
| 0.90 | 0.226 | 27 | 0.154 | -0.167 | 0.059 | 0.333 |
| 0.95 | 0.307 | 14 | 0.282 | -0.815 | 0.160 | 0.549 |

*Quantile:* data quantile used to define the threshold; *Threshold:* value of the variable at the chosen quantile; *Exceedances:* number of observations above the threshold; *Scale:* estimated GPD scale parameter ( $\beta$ ); *Shape:* estimated GPD shape parameter ( $\xi$ ); *SE Scale:* standard error of the scale parameter; *SE Shape:* standard error of the shape parameter

The Hill estimator supported this interpretation, showing a stable plateau over 26 consecutive order statistics (**Table 11**), with a highly precise average tail index of γ = 0.84 ± 0.03 (bootstrap SE). Both the plateau stability and the low coefficient of variation (5.4%) confirmed the robustness and reliability of this estimate. The tail index remaining clearly below 1 indicates moderate tail weight, providing strong support for the use of a GED model rather than a heavy-tailed (Pareto-like) distribution.

**Table 11.** Hill estimator plateau analysis for the Mataral dataset.

| Plateau k | $\gamma$ (plateau) | Hill mean | Plateau length | CV | Bootstrap SEe |
| --- | --- | --- | --- | --- | --- |
| 36 | 0.840 | 0.568 | 26 | 0.054 | 0.03 |

*Plateau k* = order statistic at plateau; $\gamma$ = tail index at plateau; *Hill mean* = mean $\gamma$ over the plateau; *Length* = number of consecutive points forming the plateau; *CV* = coefficient of variation of plateau $\gamma$ ; *Bootstrap SE* = bootstrap standard error of $\gamma$

The Generalized Extreme Value (GEV) model fitted to block maxima (**Table 12**) yielded a shape parameter close to zero (ξ = 0.008), indicating that the extreme values followed a nearly light-tailed distribution.

**Table 12.** Estimated GEV parameters for block maxima (block size 5), derived from the Mataral dataset. Columns indicate the number of blocks used, block size, the location, scale, and shape parameters, as well as the 95 % confidence intervals obtained via bootstrap (lower and upper values).

| Blocks | Block size | Location parameter | Scale parameter | Shape parameter | Location IC95 | Scale IC95 | Shape IC95 |
| --- | --- | --- | --- | --- | --- | --- | --- |
| 52 | 5 | 0.1061 | 0.1361 | 0.0083 | 0.0690<br>0.1486 | 0.0996<br>0.1667 | -0.1202<br>0.2243 |

The confidence intervals were calculated using bootstrap resampling (1000 iterations) to account for the inability to estimate asymptotic standard errors for this dataset. The bootstrap intervals provide a robust estimate of parameter uncertainty.

Tail conditional expectations (TCE) increased moderately with quantile (**Table 13**), from 0.357 at Q₀․₉ to 0.588 at Q₀․₉₉, suggesting that even the most extreme observations remained within a relatively narrow extension of the main data range.

**Table 13.** Tail Conditional Expectations (TCE) at selected quantiles (Q₀.₉, Q₀.₉₅, Q₀.₉₉) for the Mataral dataset, calculated from the GEV model fitted to block maxima. TCE values represent the expected magnitude of observations exceeding each quantile, summarizing the behavior of extreme values relative to the main distribution.

| Quantile | TCE value |
| --- | --- |
| 0.90 | 0.357 |
| 0.95 | 0.449 |
| 0.99 | 0.588 |

Estimates from the Pickands and moment methods (Supplementary Table S2) were consistent with these results: both statistics converged toward values between 0.4 and 0.6 for orders above 20, indicating a stable, moderately light tail.

### The El Chaco site

#### Basic characteristics of the distribution

**Table 3** presents the descriptive statistics of the El Chaco dataset, summarizing the main features and variability of each variable (raw data and size-corrected transformed variable) prior to further analysis. For this data set (*n*=297) and after transformation, the size-corrected variable for filament lengths ranged from –0.168 to 0.588 (standardized scale). In contrast, the raw variable ranged from 0.738 mm to 1.444 mm (range = 0.706), with a mean of 1.034 ± 0.116 and a median of 1.026, confirming its compact and symmetric structure prior to transformation. The ratio between the maximum and minimum raw values was 1.96, indicating that the longest filaments were about twice as long as the shortest ones, suggesting a relatively low variability in filament length among individuals. The transformed variable at El Chaco was characterized by a mean value close to zero (mean = –5.46 × 10⁻¹⁸), a variance of 0.013, and a standard deviation of 0.114. Because the transformation centered the data around zero, both positive and negative values were generated, rendering the coefficient of variation (CV ≈ –2.1 × 10¹⁶) meaningless. The median (–0.010) was very close to the mean, indicating only weak asymmetry. The skewness (0.67) and kurtosis (0.99) revealed a slight elongation toward the positive tail and a moderately peaked shape (**Figure 9**). The transformed variable spanned a range of 0.713, from –0.293 to 0.420, with an interquartile range (IQR = 0.151) and a median absolute deviation (*MAD* = 0.078), indicating that residual values fluctuated within narrow but balanced limits around zero.

**Figure 9.**
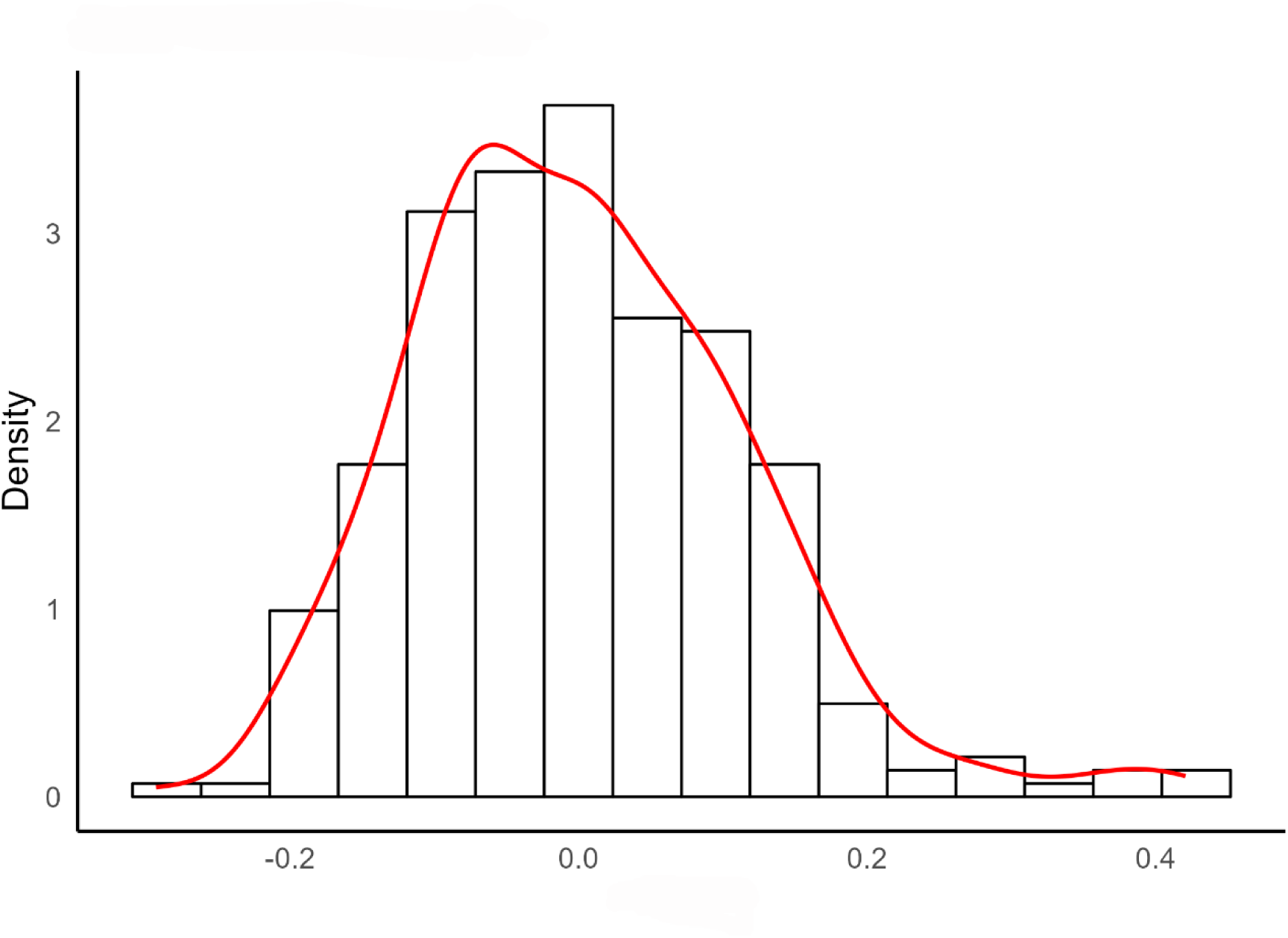
Histogram and density function of the El Chaco dataset. The variable is the size-corrected variable and represents the allometric residuals (LM-transformed mean of right and left filament lengths relative to head-collar length)

#### Outlier detection

The detection of outliers revealed only a limited number of anomalous values, with results remaining consistent across methods (**Table 4**). Between four and five observations (1.3–1.7% of the data) were detected by distance-based methods such as IQR, Z-score, and Hampel filters. Robust estimators, including the median absolute deviation (MAD), the Median of the Coupled Differences (MedCouple, a robust skewness measure), and the generalized error distribution (GED), identified between five and ten outliers (1.7–3.4% of the data). The iterative Grubbs test isolated only two points (0.7%), while no extreme values were detected by the Extreme Studentized Deviate (ESD) method. The two-component mixture model identified up to fifteen outliers (5.1%), representing the most permissive detection scenario. Overall, the dataset appeared remarkably homogeneous. The small proportion of mild outliers suggested the presence of slightly heavier tails rather than true anomalies. No evidence of extreme short filaments or distant isolated values was observed, confirming that the residual structure remained continuous and compact. **Figure 10** illustrates the distribution of observations alongside IQR-identified outliers and the 95% and 99% GED thresholds. Only a few points slightly exceed the upper limits, reflecting mild tail extensions rather than distinct clusters or extreme anomalies.

**Figure 10.**
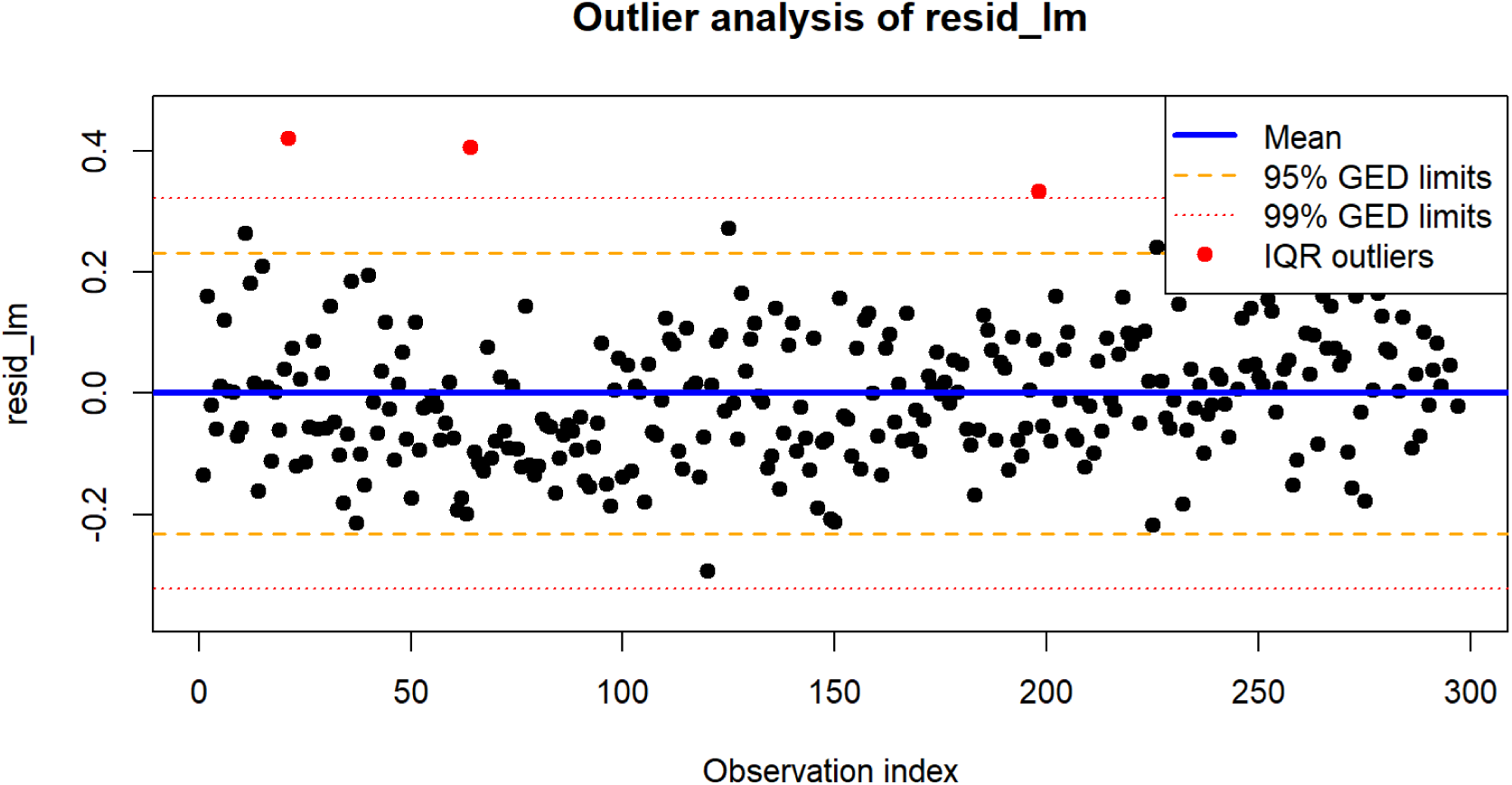
Distribution of individual data points from the Mataral dataset plotted by index (X-axis) and size-corrected variable (Y-axis). The solid horizontal line indicates the overall mean, while dashed lines represent the 95% and 99% GED (Generalized Error Distribution) limits. Red points denote outliers identified using the interquartile range (IQR) method

#### Distribution model fitting

**Table 14** presents the probability distributions evaluated for the El Chaco dataset, showing simple and mixture models. Goodness-of-fit analyses indicated that the Generalized Error Distribution (GED) provided the most accurate description of the transformed data. This model achieved the lowest AIC (–625.5) and BIC (–614.4) values, outperforming all alternative fits. The estimated GED parameters (shape *ν* = 4.23, scale *σ* = 0.26) indicated a distribution slightly more leptokurtic than the Gaussian, with moderately heavy tails but no multimodal features. The skew-normal and skew-*t* distributions ranked second and third (ΔAIC = 166.3 and 167.5, respectively), suggesting mild asymmetry but substantially lower performance. Symmetric Gaussian and Student-*t* models were less adequate, confirming that the distribution deviated from strict normality.

**Table 14.** Comparison of statistical models fitted to the filament length residuals in *An. pseudopunctipennis* larvae from El Chaco. Log-likelihood, Akaike Information Criterion (AIC), Bayesian Information Criterion (BIC), the differences relative to the best AIC (ΔAIC) and BIC (ΔBIC), and the ranking of each model based on ΔAIC and ΔBIC

| Model | Log Likelihood | AIC | BIC | $\Delta\text{AIC}$ | $\Delta\text{BIC}$ | Rank by $\Delta\text{AIC}$ | Rank by $\Delta\text{BIC}$ |
| --- | --- | --- | --- | --- | --- | --- | --- |
| GED | 315.76 | -625.52 | -614.44 | 0 | 0 | 1 | 1 |
| Skew Normal | 232.60 | -459.21 | -448.13 | 166.32 | 166.32 | 2 | 2 |
| GMMfree ($k=2$) | 234.38 | -458.75 | -440.29 | 166.77 | 174.16 | 3 | 4 |
| GMMseeded ($k=2$) | 234.38 | -458.75 | -440.29 | 166.77 | 174.16 | 3 | 4 |
| Battacharya | 234.38 | -458.75 | -440.29 | 166.77 | 174.16 | 3 | 4 |
| Skew $t$ | 233.03 | -458.05 | -443.28 | 167.47 | 171.16 | 6 | 3 |
| $t$- Student | 226.70 | -447.40 | -436.32 | 178.13 | 178.13 | 7 | 7 |
| Mixture Skew Normal ($k=2$) | 228.33 | -446.66 | -428.19 | 178.86 | 186.25 | 8 | 9 |
| Normal | 223.80 | -443.60 | -436.22 | 181.92 | 178.23 | 9 | 8 |
| Mixture t-Student ($k=2$) | 218.05 | -422.09 | -396.24 | 203.43 | 218.21 | 10 | 11 |
| Weibull | 212.20 | -420.41 | -413.02 | 205.12 | 201.42 | 11 | 10 |
| Gamma | 188.93 | -373.87 | -366.48 | 251.65 | 247.96 | 12 | 12 |
| Log-Normal | 32.97 | -61.93 | -54.54 | 563.59 | 559.90 | 13 | 13 |
| Pareto | -3578.80 | 7161.59 | 7168.98 | 7787.11 | 7783.42 | 14 | 14 |

GED: Generalized Error Distribution. GMM: Gaussian Mixture Models. All mixture models were fitted as 2 components mixtures ( $k=2$ ). The Battacharya model found 2 components

Among the mixture models, the two-component Gaussian mixture (GMM*_f_*_ree_, GMM_seeded_, Battacharya) reached the best compromise but remained inferior to the GED model (ΔAIC ≈ 166.8).

#### Two component GMM specific analysis

##### Model fitting

As with the Mataral dataset, the Bhattacharya method had difficulty clearly identifying the second component in this dataset, as it was very small and only marginally distinguishable (**Figure 11**).

**Figure 11.**
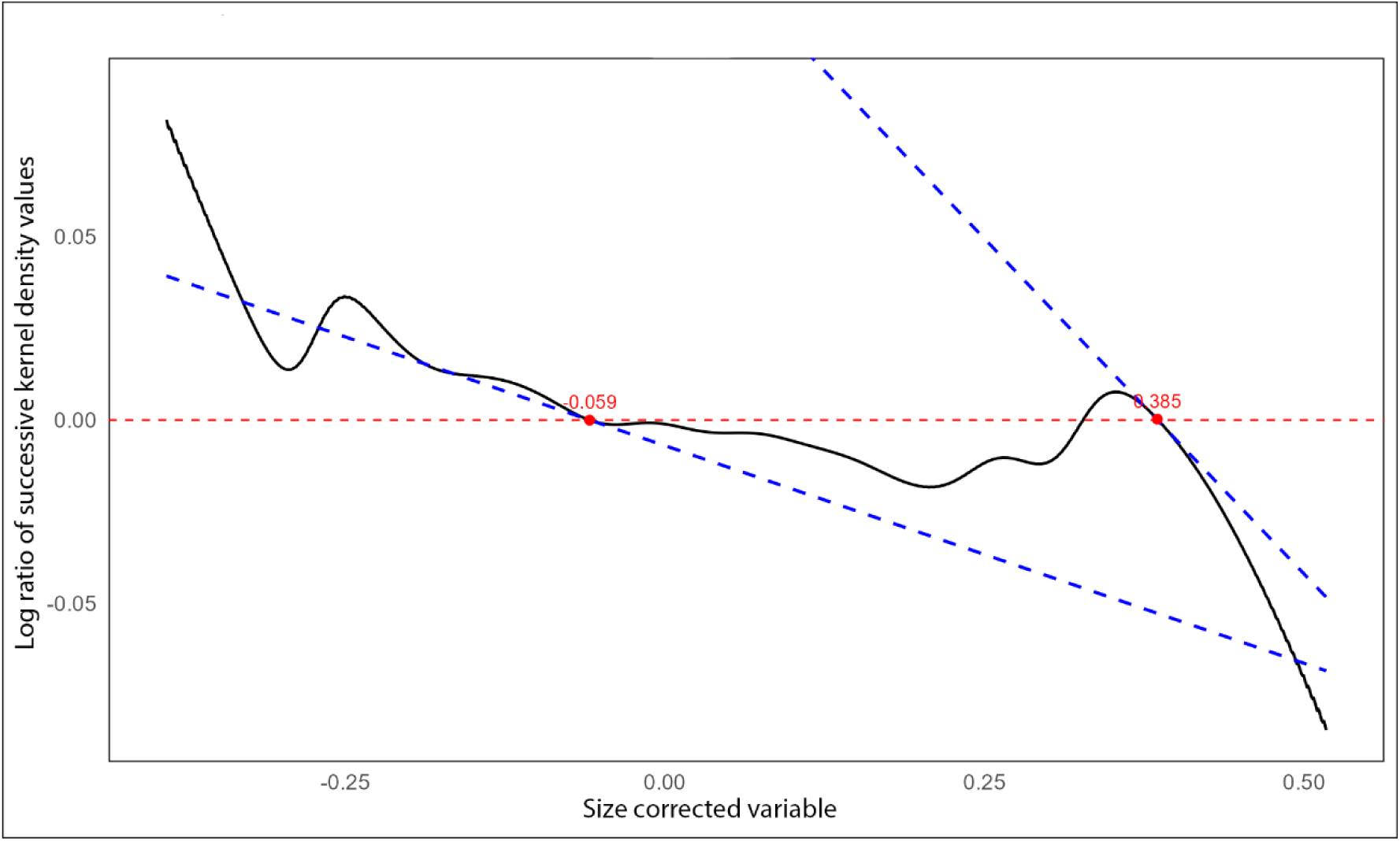
Bhattacharya plot of the EL Chaco dataset. The x-axis represents the size-corrected variable for filament lengths, and the y-axis shows the log ratio of successive kernel density values (Bhattacharya transformation). Approximate linear segments reveal the presence of two Gaussian components, from which means, variances, and proportions were estimated

Both approaches (GMM_free_ and GMM_seeded_) converged on very similar parameter estimates for the two components (**Table 15**). The major component (component 1) accounted for approximately 98.4% of the observations, with a mean near zero and a variance of ≍0.011, whereas the minor component (component 2) comprised only ≍1.6% of the data, with a mean around 0.385 and a very small variance (≍0.00082). Model likelihoods and information criteria were virtually identical across models (**Table 14**, **Table 15**). These values also confirm that, although a GMM provides a reasonable approximation, alternative distributions such as the Generalized Error Distribution (GED) may yield better fits.

**Table 15.** Summary of parameter estimates for the El Chaco dataset obtained using three estimation approaches: Bhattacharya method, Gaussian mixture model (GMM) initialized with Bhattacharya estimates (GMM*seeded*), and freely initialized GMM (GMM*free*). For each component, mean, variance, and mixing proportion (λ) are reported, along with bootstrap standard deviations of the mean (σ₍μ,boot₎) and mixing proportion (σ₍λ,boot₎)

| Model | Component | Mean | Variance | Proportion | $\sigma_{\mu,boot}$ | $\sigma_{\lambda,boot}$ |
| --- | --- | --- | --- | --- | --- | --- |
| Battacharya | 1 | -0.0061 | 0.0108 | 0.98 |  |  |
| Battacharya | 2 | 0.3846 | 0.0008 | 0.02 |  |  |
| GMMseeded | 1 | -0.0061 | 0.0108 | 0.98 | 0.0207 | 0.1841 |
| GMMseeded | 2 | 0.3846 | 0.0008 | 0.02 | 0.1108 | 0.1841 |
| GMMfree | 1 | -0.0061 | 0.0108 | 0.98 | 0.0239 | 0.2247 |
| GMMfree | 2 | 0.3846 | 0.0008 | 0.02 | 0.1133 | 0.2247 |

$\sigma_{\mu,boot}$ : Bootstrap standard deviation of the mean estimate — a measure of uncertainty on the mean; $\sigma_{\lambda,boot}$ : Bootstrap standard deviation of the mixing proportion ( $\lambda$ ) — a measure of how stable that component’s relative weight is across bootstrap samples (10 000 bootstrap resamples)

#### Component stability and separation

Bootstrapped stability analysis highlighted that the major component (component 1) was consistently stable across both GMM_free_ and GMM_seeded_, with low variability in mean and variance estimates (**Table 15**). In contrast, the minor component (component 2) displayed high bootstrap variability and was consistently classified as unstable or too small with higher relative bootstrap standard deviations, indicating low stability (**Table 15**). This instability was observed regardless of whether the GMM was freely fitted (GMM_free_) or constrained to Bhattacharyya-initialized seeds (GMM_seeded_).

Distance-based metrics confirmed the similarity of the components across models. Bhattacharyya and Mahalanobis distances between the major and minor components were identical for both models and were substantial (3.63 and 5.13, respectively) (**Table 16**), indicating a clear separation. Overlap (OVL) values were both minimal (OVL_unweighted_ ≍0.01 and OVL_weighted_ by component proportion ≍0.001) (**Table 16**), suggesting that the minor component does not substantially mix with the dominant one, but highlighting the marginal size of the minor component.

**Table 16.** Component separation and overlap for the El Chaco dataset, with distances (Bhattacharyya, Mahalanobis) and OVL values indicating the distinctness of component distributions.

| Model | Bhattacharyya distance | Mahalanobis distance | OVLunweighted | OVLweighted |
| --- | --- | --- | --- | --- |
| Batttacharya | 3.63 | 5.13 | 3.63 | 5.13 |
| GMMfree | 3.63 | 5.13 | 3.63 | 5.13 |
| GMMseeded | 3.63 | 5.13 | 3.63 | 5.13 |

#### Cross-model concordance

Cross-model stability was perfect, with 100% concordance and correlation of component probabilities equal to 1 (**Table 17**). All three models demonstrated complete concordance in point assignments, confirming that the same set of extreme values were consistently identified as belonging to component 2 (**Figure 12**).

**Figure 12.**
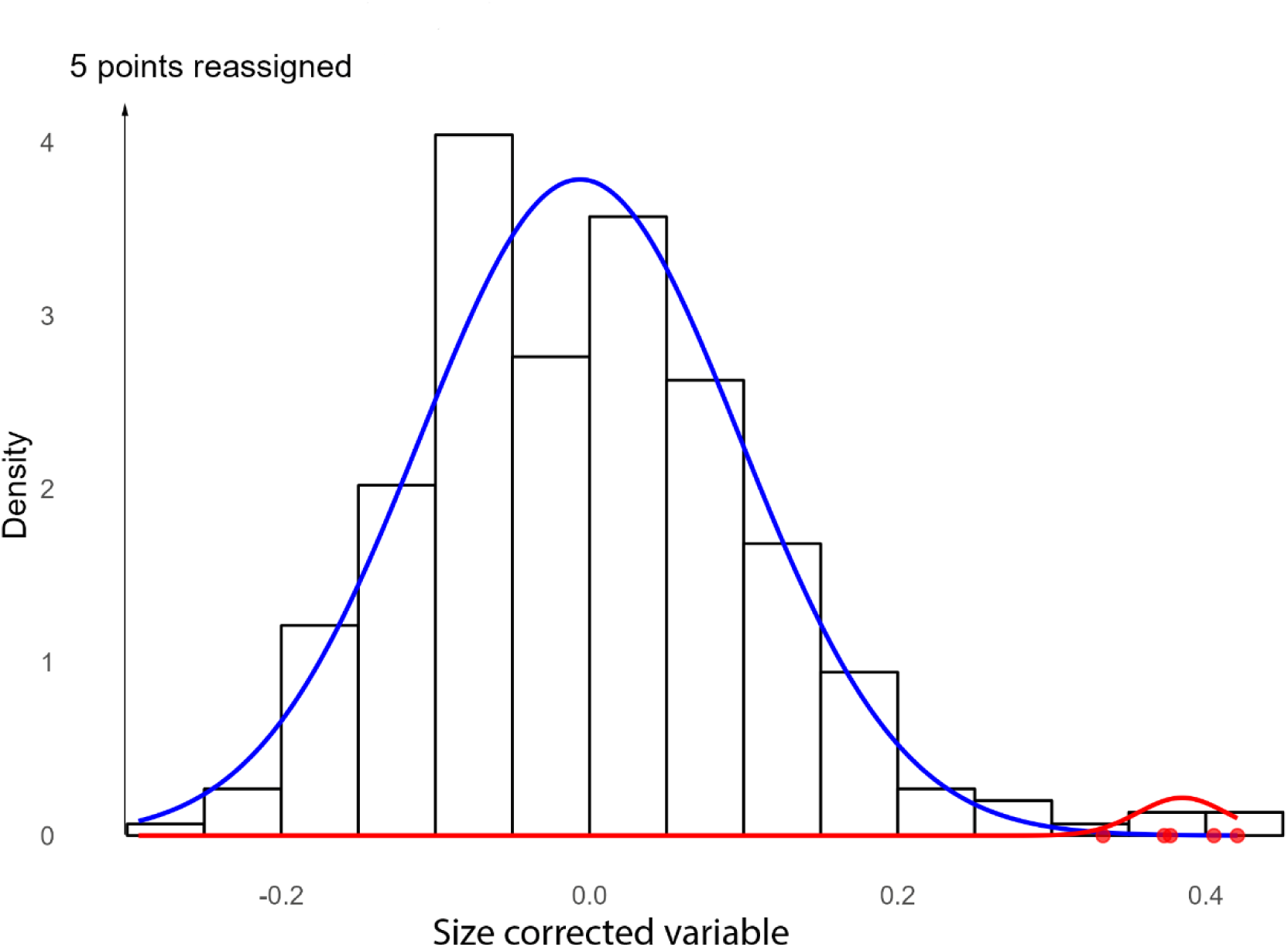
Histogram of the El Chaco dataset showing fitted Gaussian components using the Bhattacharya method, GMM*free*, and GMM*seeded* approaches (all three methods identify the same components), with data points reassigned according to posterior probabilities

**Table 17.** Concordance of component assignments between models for the Mataral dataset, with percentage agreement and correlations of membership probabilities indicating similarity of clustering.

| Model 1 | Model 2 | Concordance | Corr-p1 | Corr-p2 |
| --- | --- | --- | --- | --- |
| Battacharya | *GMMfree* | 100 | 1 | 1 |
| Battacharya | *GMMseeded* | 100 | 1 | 1 |
| *GMMfree* | *GMMseeded* | 100 | 1 | 1 |

Inspection of individual reassigned points revealed that only a very small fraction of the dataset had high probability of belonging to component 2 (Supplementary Table S3), consistent with the low proportion of this component. For example, the highest-valued points (>0.33) exhibited a high probability of assignment to the minor component (>0.7–0.99), further illustrating that component 2 represents the extreme tail of the distribution rather than a substantial subpopulation.

Overall, these results indicate that while the data can be formally represented as a mixture of two Gaussians, the minor component is extremely small and unstable. This suggests that component 2 likely reflects the extreme values of the main population rather than a true biological subpopulation. The major component encompasses the vast majority of the data, with the few points assigned to the minor component located at the upper extreme of the distribution.

#### Tail specific analysis

##### Multimodality and subpopulation tests

Tests of multimodality consistently supported unimodality. The Dip test (*p* = 0.676), Silverman test (*p* = 0.414), and Excess Mass test (*p* = 0.211) all failed to reject the null hypothesis of a single mode. The Mode Tree analysis confirmed one dominant mode, and model-based clustering (ΔBIC = 2) supported a single component solution. However, the Gap statistic indicated a local optimum at *K* = 3 (Gap = 0.113 ± 0.096), suggesting weak internal partitioning possibly caused by tail elongation rather than true multimodality (**Table 18**).

**Table 18.** Gap statistic and standard error (SE) for different numbers of clusters in the El Chaco dataset.

| K<br>(number of<br>clusters) | Gap value | Standard error<br>(SE) |
| --- | --- | --- |
| 1 | 0.037 | 0.099 |
| 2 | 0.060 | 0.085 |
| 3 | 0.059 | 0.115 |

The detection of 15 large gaps in the consecutive sorted data corroborates the presence of sparse but distinct high-end values, although not forming a separate mode. Altogether, these results indicated a unimodal structure with limited subpopulation differentiation.

#### Characterization of the right-hand tail

The right-hand tail was found to be moderately heavy, with a small number of extreme values contributing to its elongation. At the 0.9 quantile (threshold = 0.142), the Generalized Pareto Distribution (GPD) yielded a positive shape parameter (ξ = 0.167 ± 0.30), indicative of a weakly heavy tail. Beyond the 0.95 quantile (threshold = 0.182), the shape parameter became negative (ξ = –1.03 ± 0.002), suggesting that the upper tail was bounded (**Table 19**). This bounded behavior was consistent with the limited amplitude observed in the raw and transformed variables.

**Table 19.** Generalized Pareto Distribution (GPD) fits to the upper tail of the El Chaco dataset.

| Quantile | Threshold | Exceedances | Scale | Shape | SE Scale | SE Shape |
| --- | --- | --- | --- | --- | --- | --- |
| 0.90 | 0.142 | 30 | 0.065 | 0.167 | 0.023 | 0.299 |
| 0.95 | 0.182 | 15 | 0.245 | -1.030 |  | 0.002 |

*Quantile:* data quantile used to define the threshold; *Threshold:* value of the variable at the chosen quantile; *Exceedances:* number of observations above the threshold; *Scale:* estimated GPD scale parameter ( $\beta$ ); *Shape:* estimated GPD shape parameter ( $\xi$ ); *SE Scale:* standard error of the scale parameter; *SE Shape:* standard error of the shape parameter

The Hill estimator (γ = 0.512 ± 0.229) did not exhibit a stable plateau, indicating limited robustness of tail estimates and supporting a finite-tail model rather than a power-law decay (**Table 20**).

**Table 20.** Hill estimator plateau analysis for the El Chaco dataset.

| Plateau k | $\gamma$ (plateau) | Hill mean | Plateau length | CV | Bootstrap SEe |
| --- | --- | --- | --- | --- | --- |
| 76 | 1.799 | 0.512 | 8 | 0.046 | 0.023 |

*Plateau k* = order statistic at plateau; $\gamma$ = tail index at plateau; *Hill mean* = mean $\gamma$ over the plateau; *Length* = number of consecutive points forming the plateau; *CV* = coefficient of variation of plateau $\gamma$ ; *Bootstrap SE* = bootstrap standard error of $\gamma$

The Generalized Extreme Value (GEV) fit provided a location of 0.091, a scale of 0.087, and a negative shape parameter (–0.104), pointing to a Weibull-type tail with finite support (**Table 21**)

**Table 21.** Estimated GEV parameters for block maxima (block size 5), derived from the El Chaco dataset. Columns indicate the number of blocks used, block size, the location, scale, and shape parameters, as well as the 95 % confidence intervals obtained via bootstrap (lower and upper values).

| Blocks | Block size | Location parameter | Scale parameter | Shape parameter | Location IC95 | Scale IC95 | Shape IC95 |
| --- | --- | --- | --- | --- | --- | --- | --- |
| 59 | 5 | 0.0914 | 0.0867 | -0.1042 | 0.0701 | 0.0640 | -0.2317 |
|  |  |  |  |  | 0.1126 | 0.1027 | 0.0526 |

The confidence intervals were calculated using bootstrap resampling (1000 iterations) to account for the inability to estimate asymptotic standard errors for this dataset. The bootstrap intervals provide a robust estimate of parameter uncertainty.

Conditional tail expectations (TCE) increased smoothly from 0.219 at Q_0_.₉ to 0.401 at Q₀.₉₉, illustrating a moderate but bounded growth of extremes (**Table 22**)

**Table 22.** Tail Conditional Expectations (TCE) at selected quantiles (Q₀.₉, Q₀.₉₅, Q₀.₉₉) for the El Chaco dataset, calculated from the GEV model fitted to block maxima. TCE values represent the expected magnitude of observations exceeding each quantile, summarizing the behavior of extreme values relative to the main distribution.

| Quantile | TCE value |
| --- | --- |
| 0.90 | 0.219 |
| 0.95 | 0.279 |
| 0.99 | 0.401 |

The Pickands and Moment estimators fluctuated between 0.2 and 1.8 across orders, without stabilization, further confirming the interpretation of a light to moderately heavy but finite tail (Supplementary Table S4).

Altogether, these results suggest that the apparent “extremes” are simply large individuals arising from the same quasi-Gaussian distribution, rather than representing a separate group of outliers.

## Discussion

Continuous biological traits such as body size, length, or weight are often assumed to follow a normal (Gaussian) distribution, reflecting the additive influence of many small genetic and environmental effects (Fisher, 1918; Mackay, 2009; Neher & Shraiman, 2011). This expectation, rooted in the Central Limit Theorem and the infinitesimal model of quantitative genetics, has made the Gaussian distribution a standard *a priori* model in biological data analysis. However, empirical evidence shows that many traits deviate from perfect normality. For example, many continuous traits — especially those constrained to positive values— exhibit right-skewed distributions, where most individuals cluster near lower values and a few extreme individuals form a positive tail (Taylor-LaPole et al., 2022; Cao et al., 2023). In such cases, a log-normal or gamma distribution often provides a better fit to the data (Scherer et al., 2020; Díaz et al., 2022). Even morphometric traits such as plant height or organ length, which can appear approximately symmetric, may show significant skewness or kurtosis when measured across heterogeneous populations or developmental stages (Di Santo et al., 2021; Gearty et al., 2024). Thus, while the normal law remains a useful approximation, verifying distributional assumptions remains essential for biological inference. Recognizing and quantifying departures from normality also provides insight into the underlying biological processes — such as developmental constraints, allometric scaling, or selective asymmetries — that shape natural variation.

Gaussian mixture models (GMMs) are widely used in biology to identify subpopulations or complex structures within heterogeneous biological datasets (McClellan & Logan, 1994; Lardeux & Tetuanui, 1995; Wu et al., 2013; Sukovata, 2019; Nussbaumer et al., 2021). However, this approach can become challenging when small subpopulations are mixed within much larger ones, particularly in two-component mixtures, as observed in the present study. When a small number of observations occur in the tail of a distribution, the GMM may not recognize them as a separate component and may “absorb” sparse tail data into nearby, denser components, as the EM algorithm prioritizes overall likelihood. When recognized, the GMM may estimate the mean of the minor component closer to that of the main population than suggested by the Bhattacharya-based initialization. Bhattacharya-based initialization can detect minor modes in the tail, but these peaks often involve very few points and are not consistently stable across resampling. Therefore, the presence of a minor component in the tail does not necessarily indicate a biologically meaningful subpopulation, but may instead reflect tail fluctuations of the main population.

Model selection based on AIC and BIC supported the Generalized Error Distribution (GED) as providing the best fit to the present datasets. The greater flexibility of the GED allows it to account for skewness and heavy tails that are not adequately represented by a simple two-component Gaussian mixture.

*Anopheles pseudopunctipennis* has historically been divided into multiple subspecies based on morphological differences across its broad geographic range (Harbach & Wilkerson, 2023). The historical designation of multiple subspecies, despite recent taxonomic revisions, reflects the inherent phenotypic variability of the species. Morphometric analyses across populations in South America have documented significant variation in traits such as length of proboscis, length of and dark scale spots on the wing costal vein, reflecting phenotypic plasticity in response to environmental conditions (Dantur et al., 2011). Genetic studies using mitochondrial COI sequences further demonstrate high haplotype diversity and low population differentiation, suggesting extensive gene flow and the capacity to adapt to diverse ecological niches ((Dantur Juri et al., 2014). Electrophoretic surveys of multiple populations also reveal considerable variability in allele frequencies at several enzyme loci, highlighting underlying genetic heterogeneity (Manguin et al., 1995). Taken together, these findings indicate that phenotypic variation is an inherent feature of this species, and that extreme filament lengths are therefore to be expected.

In the two studied sites, El Chaco and Mataral, caudal filament lengths of larvae of *An. pseudopunctipennis* displayed a broad range, up to approximately two-fold between the smallest and largest individuals. This substantial variation can create the impression, when observing only a few individuals in field samples, that multiple subpopulations exist—a pattern consistently noted across nearly all field collections from the region (Lardeux et al., 2009). Despite this apparent heterogeneity, filament length distributions were broadly similar between the two sites under study, with minor differences in tail characteristics. In both sites, the distributions deviated from normality, and the generalized error distribution (GED) provided a superior fit, capturing moderate skewness and kurtosis typical of continuously varying traits influenced by multiple additive factors. Although a few individuals exhibited unusually long filaments, statistical analyses revealed no bimodality or distinct secondary components, indicating that these extremes represent the continuous tail of a single heterogeneous population rather than separate subgroups. The occurrence of “large filament” individuals likely reflects natural inter-individual variability, phenotypic plasticity, or stochastic growth trajectories, potentially influenced by larval nutrition, microhabitat conditions, developmental timing, or genetic variation. Their rarity and the absence of heavy tails in extreme-value analyses further support the interpretation that they are ordinary, albeit infrequent, outcomes within the population. Functionally, variation in filament length could have ecological or behavioral implications, for example influencing sensory perception or oviposition, though this remains to be tested. The observed homogeneity between sites also suggests that populations experience broadly similar environmental conditions, limiting site-specific genetic differentiation.

Methodologically, these findings highlight the importance of selecting appropriate distributional models, as assuming normality could obscure the presence of extreme phenotypes or misrepresent trait variability. While the sample sizes were sufficient to robustly characterize the overall population structure in the present study, caution is warranted in interpreting the biological drivers of the most extreme values. Future studies manipulating larval resources, assessing microclimatic variation, analyzing genotype–phenotype relationships, or tracking temporal/seasonal cohorts could clarify whether extreme filaments recur under specific conditions or arise sporadically, helping to disentangle ecological, genetic, and stochastic contributions to filament-length variation.

Understanding the natural variability and distributional extremes of morphological traits (not only the larval filament length) may provide insights into mosquito development, survival, and dispersal, with potential implications for vectorial capacity and population dynamics.

## Conclusion

Statistical analysis of filament-length distributions in *An. pseudopunctipennis* reveals continuous variation, with rare extremes forming the natural tail rather than distinct subpopulations. This pattern aligns with the species’ known morphological and genetic variability, including documented phenotypic plasticity. Distributions of filament length from the two study sites deviated from normality and were best described by a Generalized Error Distribution (GED), highlighting the importance of appropriate statistical models. While these findings characterize population-level patterns, experimental studies will be needed to determine the environmental, developmental, and genetic factors underlying extreme filament lengths and their potential ecological or vectorial consequences.

## Data availability statement

The data and R scripts that supports the findings of this study are available in theZEnodo repository: https://doi.org/10.5281/zenodo.17712350

## Data, scripts, code, and supplementary information availability

Data are available online: DOI of the webpage hosting the data https://doi.org/10.5281/zenodo.17712350, Lardeux et al. 2025.

Scripts and code are available online: DOI of the webpage hosting the data https://doi.org/10.5281/zenodo.17712350, Lardeux et al. 2025.

Supplementary information is available online: DOI of the webpage hosting the data https://doi.org/10.5281/zenodo.17712350, Lardeux et al. 2025.

